# Single Cell-type Integrative Network Modeling Identified Novel Microglial-specific Targets for the Phagosome in Alzheimer’s disease

**DOI:** 10.1101/2020.06.09.143529

**Authors:** Kruti Rajan Patel, Kuixi Zhu, Marc Y.R. Henrion, Noam D. Beckmann, Sara Moein, Melissa L. Alamprese, Mariet Allen, Xue Wang, Gail Chan, Thomas Pertel, Parham Nejad, Joseph S. Reddy, Minerva M. Carrasquillo, David A Bennett, Nilüfer Ertekin-Taner, Philip L. De Jager, Eric E. Schadt, Elizabeth M. Bradshaw, Rui Chang

## Abstract

Late-Onset Alzheimer’s Disease (LOAD) results from a complex pathological process influenced by genetic variation, aging and environment factors. Genetic susceptibility factors indicate that myeloid cells such as microglia play a significant role in the onset of LOAD. Here, we developed a computational systems biology approach to construct probabilistic causal and predictive network models of genetic regulatory programs of microglial cells under LOAD diagnosis by integrating two independent brain transcriptome and genome-wide genotype datasets from the Religious Orders Study and Rush Memory and Aging Project (ROSMAP) and Mayo Clinic (MAYO) studies in the AMP-AD consortium. From this network model, we identified and replicated novel microglial-specific master regulators predicted to modulate network states associated with LOAD. We experimentally validated three microglial master regulators (*FCER1G*, *HCK* and *LAPTM5*) in primary human microglia-like cells (MDMi) by demonstrating the molecular impact these master regulators have on modulating downstream genomic targets identified by our top-down/bottom-up method and the causal relations among the three key drivers. These master regulators are involved in phagocytosis, a process associated with LOAD. Thus, we propose three new master regulator (key driver) genes that emerged from our network analyses as robust candidates for further evaluation in LOAD therapeutic development efforts.

## Introduction

Late-Onset Alzheimer’s disease (LOAD) is a complex neurodegenerative disease that is characterized by neuropathology consisting of amyloid beta (Aß) plaques, neurofibrillary tangles and clinical dementia. Genome-wide association studies (GWAS) have implicated immune cell-specific genes associated with AD risk that point to microglia as a causal cell type [1–18]. Microglial cells are resident innate immune cells of the central nervous system[19] and play an important role in abolishing apoptotic cells, Aß deposits and synapse removal by phagocytosis. It has previously been discovered that microglia are associated with amyloid plaques in the brain[20]. Especially, given the recently failed clinical trials on anti-amyloid plaques[21], it becomes critically important to understand molecule mechanisms of microglia in the formation and clearance of amyloid plaques. A previous study using co-expression network analysis of post-mortem brains from patients with LOAD showed a microglial module dominated by genes implicated in phagocytosis[22].

In this study, we sought to identify cell type specific master regulators (key drivers) modulating network states underlying microglial functions, particularly those involved in phagocytosis and Aβ clearance in the context of AD. We applied the computational framework PSEA [23] to deconvolve bulk-tissue RNA sequencing (RNA-seq) data from post-mortem brain regions to isolate the microglial-specific gene expression signal. The reason we chose the PSEA method over other popular methods, such as Cibersort [24], dtangle [25], DSA [26] and NNLS [27], is that these methods cannot directly estimate cell-type specific residuals from the bulk-tissue RNA-seq data, instead, these methods only estimate cell fraction in a bulk-tissue sample. We demonstrated the robustness of this de-convolution method using random selection of microglial biomarkers derived from single-cell RNA-seq (scRNAseq) studies [28–32]. Next, we applied a novel systems biology approach to these data to build microglial-specific probabilistic causal network models of the immune component of AD. From these models we identified master regulators of the network states in microglial cells for AD. Among the predicted microglial-specific key drivers, we experimentally validated three novel targets, *HCK, FCER1G, and LAPTM5,* that replicated across our two cohorts using human monocyte-derived microglial-like (MDMi) cells[33]: *HCK* is a member of the Src family of protein tyrosine kinases which couples to Fc receptors during phagocytosis. *FCER1G* is predominantly expressed by hematopoietic cells, encodes for a subunit of an IgE Fc receptor and is implicated as a hub gene in amyloid overexpressing models[34]. Both *FCER1G* and *HCK* were found to be hub genes of microglial modules in a LOAD transcriptomics study[22]. A study by Castillo and colleagues[35] shows that both genes are upregulated in the cortex of *App*^NL-G-F/NL-G-F^ transgenic mice as Aβ amyloidosis progresses. *FCER1G* shows a significant association in regards to immune and microglial functions and amyloid deposits in humans and mice[34, 36]. Lastly, *LAPTM5* is associated with lysosomes organization and biogenesis[37–40]. A recent study using a murine amyloid responsive network and GWAS defined association has shown that genetic variations in *LAPTM5* are associated with amyloid deposition in AD[41].

In the present study, by using causal predictive network modeling, we showed that *HCK*, *FCER1G* and *LAPTM5* not only function together in the same co-expression module (subnetwork), but they also play a key driver role in AD through effects on phagocytosis and lysosomal function in microglial cells. In addition, our network model showed that *HCK* is downstream of both *FCER1G* and *LAPTM5*, and that *FCER1G* is downstream of *LAPTM5*. Finally, we validated these predicted relationships and their functions in the MDMi model[33]: we used lentivirus mediated targeted shRNA to knockdown *HCK*, *FCER1G* and *LAPTM5* in MDMi cells and then measured the expression of their downstream genes as predicted by our microglia-specific network model. Not only did our predictive network model accurately predict the genes that changed in response to these perturbations, but it accurately predicted the gene expression dynamics as well of the downstream genomic targets. In addition, we also confirmed a causal role for two of the key drivers as modulators of phagocytic function of microglia cells using an Aß uptake assay.

## Results

### Integrative Systems Biology Approach for Constructing Single Cell-Type Regulatory Networks of AD

We developed an integrative network analysis pipeline (Figure 1) to construct data-driven microglial-specific predictive networks of AD. The overall strategy for elucidating the single cell-type gene network model depicted in Figure 1 is centered on the objective, data-driven construction of predictive network models of AD that can then be directly queried to not only identify the network components causally associated with AD, but to identify the master regulators of these components and the impact they have on the expression dynamics of the genes comprising the biological processes underlying AD, moving us towards predictive molecular models of diseases. We previously developed and applied the network reconstruction algorithm, top-down & bottom-up predictive network (predictive network for short), which statistically infers causal relationships between DNA variation, gene expression, protein expression and clinical features that are scored in hundreds of individuals or more[42].

**Figure 1.**
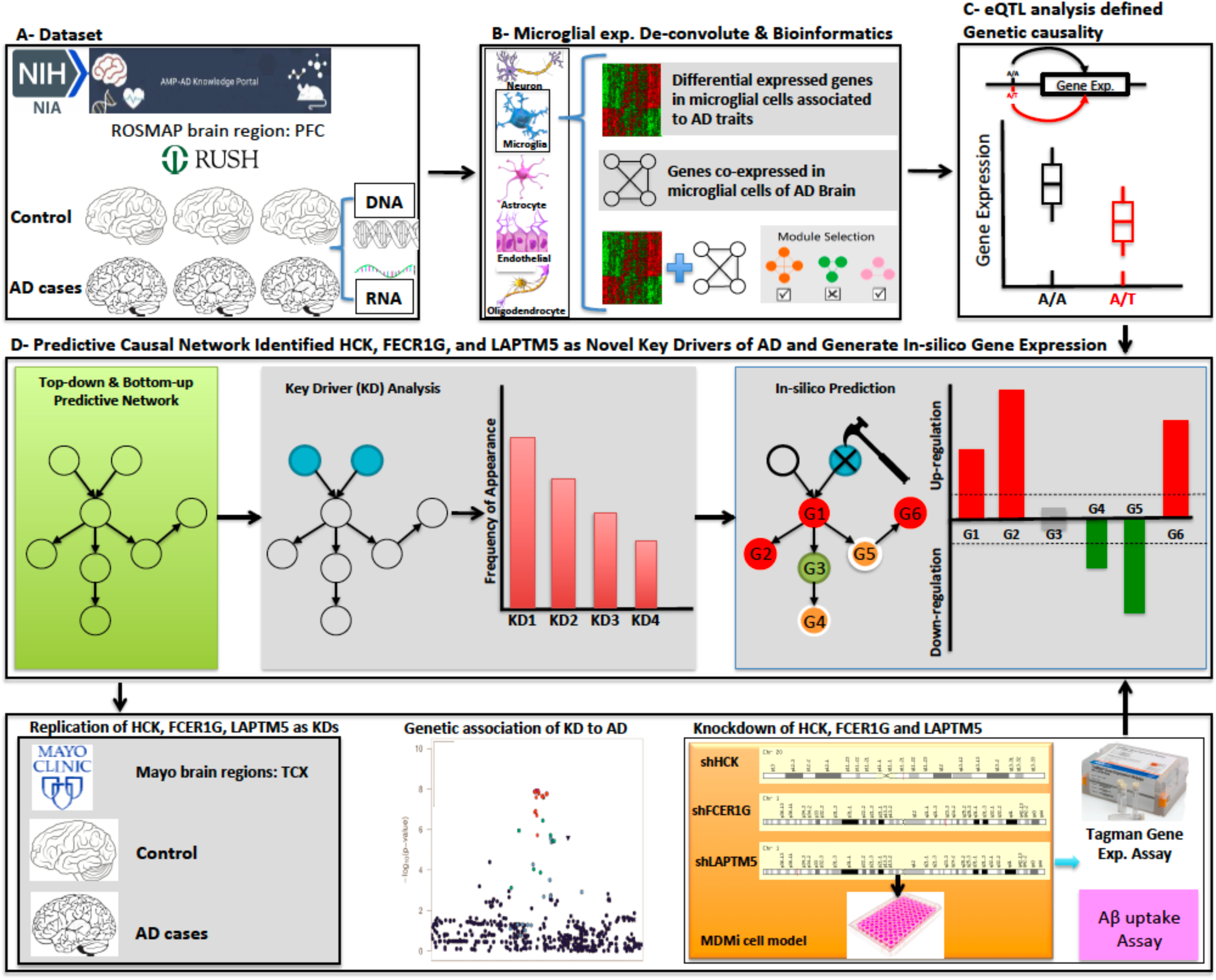

The inputs required for this type of analysis are the molecular and clinical data generated in populations of individuals, as well as first order relationships between these data, such as QTL mapped for the molecular traits and causal relationships among traits inferred by causal mediation analysis that use the mapped QTL as a source of perturbation. These relationships are input as structure priors to the network construction algorithm, boosting the power to infer causal relationships at the network level, as we and others have previously shown [22, 43–50].

To focus on the microglial component of AD, we applied the PSEA de-convolution algorithm [23] to the transcriptomic data to identify the microglial-specific expression component of the transcriptome data (Step 1, Figure S1). We demonstrated this method is robust against random selection of microglia biomarkers derived from single-cell RNA-seq (scRNAseq) studies[28–32]. Given the microglial expression component in the ROSMAP and MAYO populations, we further focused the input of molecular traits into the network reconstruction algorithm on those traits associated with AD, by identifying AD gene expression signatures comprised of hundreds to thousands of gene expression traits (Step 2, Figure S1). These signatures were enriched for a number of pathways, including mitochondrial and immune processes. To identify gene expression traits co-regulated with the AD signature genes, we constructed gene co-expression networks, and from these networks identified highly interconnected sets of co-regulated genes (modules) that were significantly enriched for the AD expression signatures as well as for pathways previously implicated in AD (Step 3, Figure S1). To obtain a final set of genes for input into the causal network construction process, we combined genes in the co-expression network modules enriched for AD signatures (the seed set, Step 5, Figure S1).

With our AD-centered input set of microglial genes for the network constructions defined, we mapped expression quantitative trait loci (eQTLs) for microglial-specific gene expression traits to incorporate the QTL as structure priors in the network reconstructions, given they provide a systematic perturbation source that can boost the power to infer causal relationships (Step 4, Figure S1)[22, 43, 44, 46–51]. The input gene set, and eQTL data from ROSMAP were then processed by the predictive network to construct probabilistic causal networks of AD (Step 6, Figure S1). An artificial intelligence algorithm to detect key driver genes from these network structures was then applied to identify and prioritize master regulators of the AD networks (Step 7, Figure S1). An in-silico prediction algorithm was developed and used to predict expression profiles upon perturbation. Our findings were then replicated in the MAYO dataset. For the top regulators we identified, we performed functional and molecular validation in microglial cell systems.

### The ROSMAP/Mayo Clinic Study Populations and Data Processing

Our predictive network pipeline starts by integrating whole exome sequencing (WES) and RNA sequencing (RNA-seq) data generated from the dorsolateral prefrontal cortex of 612 persons from ROSMAP[52–55] and from the temporal cortex of 266 patients from MAYO [56–58] in the Accelerating Medicines Partnership - Alzheimer’s Disease (AMP-AD) consortium, spanning the complete spectrum of AD clinical and neuropathological traits (Figure 1). We processed matched genotype and RNA-seq data (Online Methods). CNS tissue consists of various cell types, including neurons, endothelial and glial cells. To discover key network drivers, which could serve as therapeutic targets, specific to a single cell type in the CNS and study their contribution to AD in that specific cell type, we utilized well-known/verified single-cell marker genes to directly de-convolve bulk-tissue gene expression data into cell type-specific gene expression for the five primary cell types in the CNS: neurons, microglia, astrocytes, endothelia and oligodendrocytes (Online Methods). In this study, we focused on investigating the role of microglial cells in AD due to their strong genetic association with AD pathogenesis [59–61]. After normalizing RNA-seq data, we evaluated the contribution of demographic, clinical, technical covariates and cell-specific markers to the gene expression variance of the 5 primary cell types using a variance partition analysis (VPA)[62] (Figure 2A). The list of cell type-specific marker genes used for neurons, microglia, astrocytes, endothelia and oligodendrocytes were *ENO2*, *CD68*, CD34, *GFAP*, and *OLIG2* respectively, as previously published[57].

**Figure 2.**
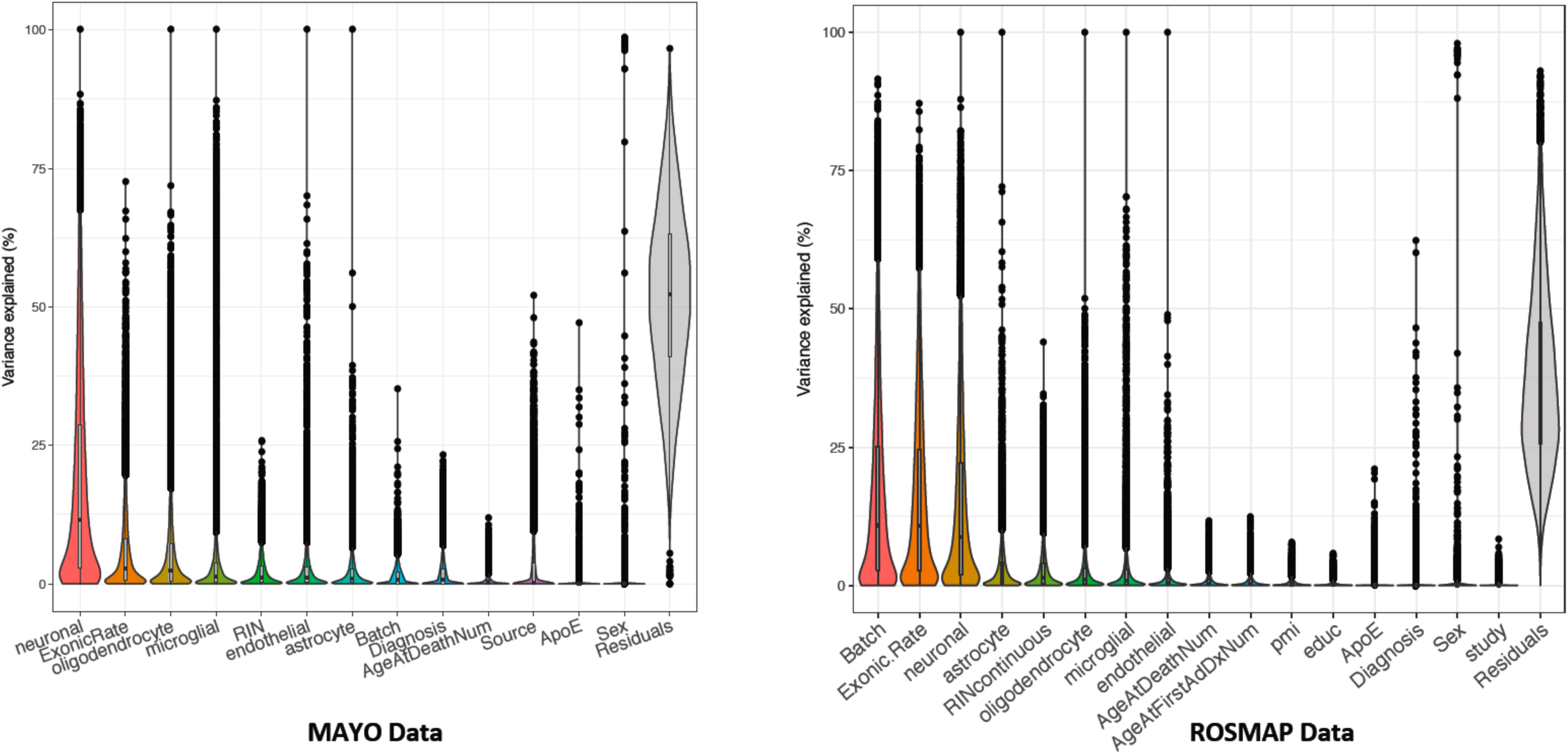
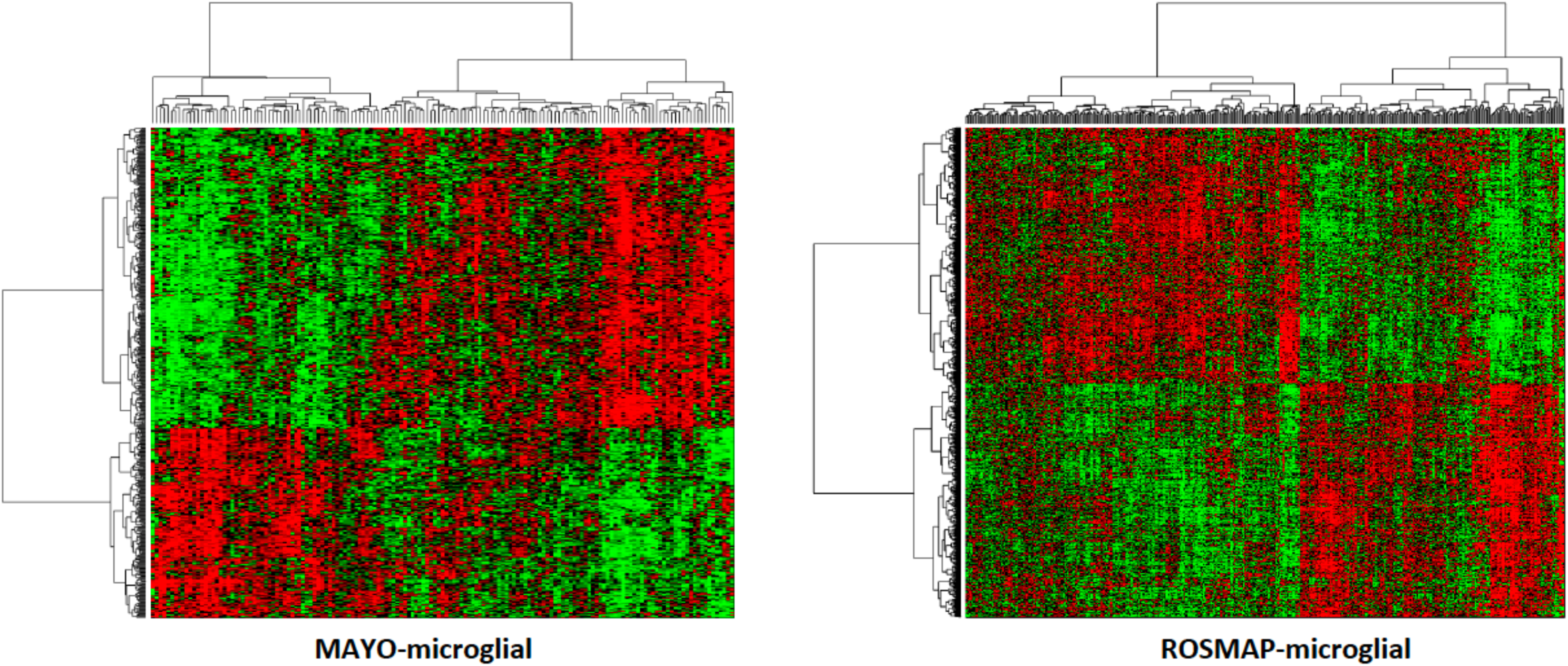
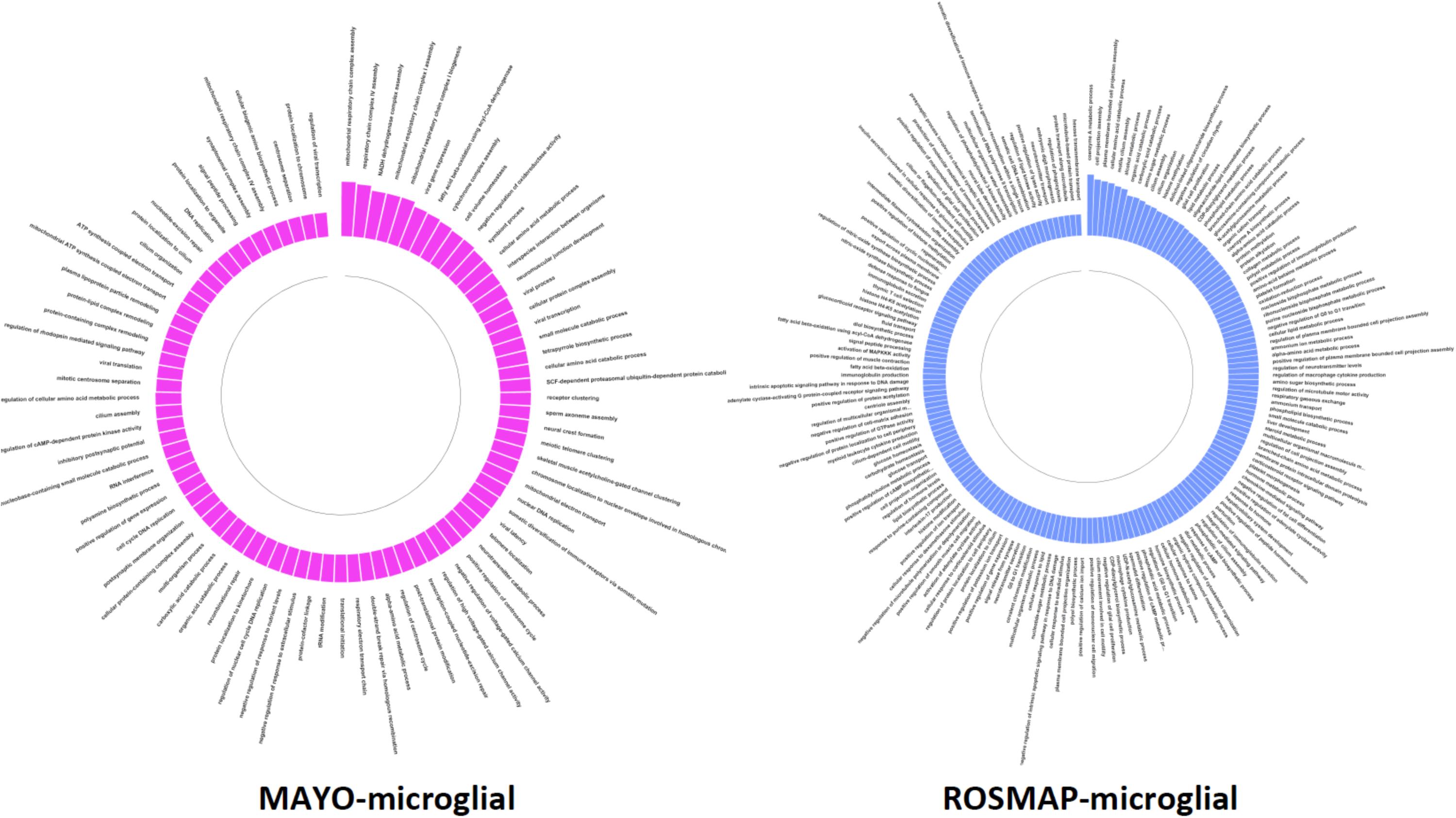

The rational of using single-gene biomarkers in the above over multi-gene biomarkers derived from scRNAseq data is as follows: 1) multi-gene biomarkers derived from various scRNAseq studies in human brain under control conditions [28–32] shows non-significant (FDR>0.05, Online Method) overlap (Figure S2, 1/0/1/1 significant pair out of 6 study-pairs in Asctrocyte/Endothelial/Microglial/Oligodendrocyte types and 2 significant pairs out of 10 study-pairs in Neuron), indicating lack of robustness and consensus in these biomarkers derived from these studies; 2) significant overlap of scRNAseq-derived biomarkers expression in ROSMAP and MAYO AD samples by PCA analysis (Figure S3), indicating the majority of scRNAseq-derived biomarkers are not ideal in distinguishing cell populations under AD conditions; 3) Significant overlap between scRNAseq-derived biomarkers and AD therapeutic targets (Figure S4) in the AMP-AD AGORA knowledge portal (https://agora.ampadportal.org/genes). This overlap is more significant than randomly selected genes from the background overlapping with the AGORA list (Online method), indicating that scRNAseq-derived biomarkers may play a significant role in AD pathology. Therefore, they are not optimal for adjusting the bulk-tissue gene expression variance by PSEA. By contrast, our single-gene biomarker, though not perfect, is derived from biological knowledge and validated by others [57]. Moreover, our single-gene biomarker had no overlap with AD therapeutic targets in AGORA, which make them good candidates for PSEA. 4) Further, our single-gene biomarker derived microglial-specific residual is significantly (p-value<2.2E-16, Figure S5, Online Method) correlated with the “pseudo” microglial-specific residuals derived from randomly selected subset of scRNAseq biomarkers by PSEA, indicating that our single-gene biomarker derived microglial-specific residual represents a robust microglial component in the bulk-tissue RNAseq data for following analysis.

In addition to the cell-type specific markers, in ROSMAP, the covariates used in VPA included Sequencing Batch, Exonic Mapping Rate, RNA Integrity Number (RIN), Age-at-death, Age-at-first-AD-diagnosis, Post-Mortem Interval (PMI), Education, *APOE* genotype, Diagnosis, Sex and Study. In MAYO, we included Exonic Mapping Rate, RIN, Sequencing Batch, Diagnosis, Age-at-death, Tissue source, *APOE* genotype and Sex. Next, cell-type specific gene expression residuals were calculated by adjusting the bulk tissue expression data by these covariates and the cell-type specific markers using a linear regression model. To get cell-type specific gene expression, including microglial-specific gene expression, we added the estimated variance of each cell type to the residual (Online Methods). In this way, the cell-type specific gene expression can be directly derived from expression data without the need to first estimate the cell population from bulk tissue data, which could induce approximation errors.

### Identifying an AD-associated gene set in microglia and mapping their eQTL

To identify an AD-centered set of microglia gene expression traits, we performed microglial-specific differential expression (DE) analysis in the ROSMAP and MAYO cohorts using the derived microglial-specific expression data (Online Methods). By comparing expression data from AD and pathologically confirmed controls (CN), there were 513 significantly (FDR<0.05) DE microglial-specific genes in the MAYO dataset (MAYO-microglial for short) and 1,693 microglial-specific genes in the ROSMAP dataset (ROSMAP-microglial for short) (Figure 2B), with 120 significant DE genes overlapping between these two sets (Fisher Exact Test, odd ratio=3.8169, p-value<2.2E-16). To examine the biological processes that are disrupted between AD cases and controls, as reflected in the DE signatures, we performed Pathway Enrichment Analysis (PEA) on the DE gene set for each cohort. We identified 84 and 173 GO terms (Figure 2C), and 15 and 54 KEGG pathways (Figure S6) that are enriched (p-value<0.05) in the MAYO and ROSMAP-microglial data respectively. These pathways include well-known biological functions associated to AD, such as mitochondrial functions, amino acid metabolism, lipid metabolism, glial cell functions, Fc gamma R-mediated phagocytosis, Phagosome etc.

Another critical input for the construction of predictive network models, are the expression quantitative trait loci (eQTL), leveraged as a systematic source of perturbation to enhance causal inference among molecular traits, an approach we and others have demonstrated across a broad range of diseases and data types[22, 43–47, 49–51, 63–77]. We mapped cis-eQTL by examining the association of microglial-specific expression traits with genome-wide genotypes [78–82] assayed in the ROSMAP and MAYO cohorts (Online Methods). In the MAYO-microglial, 3875 (19.5%) of the genes tested were significantly correlated with allele dosage (FDR < 0.01) and in ROSMAP-microglial, 5186 (25.6%) of the genes tested were significantly correlated with allele dosage (FDR < 0.01). Of the cis-eQTL detected in each cohort, 1785 eQTL (46% in MAYO and 34% in ROSMAP) were overlapping between the two sets.

### Microglial-specific Co-expression Network in AD

While DE analysis can reveal patterns of microglial-specific expression associated with AD, the power of such analysis to detect a small-to-moderate expression difference is small. To complement the DE analyses in identifying the input gene set for the predictive network, we clustered the microglial gene expression traits into data-driven, coherent biological pathways by constructing co-expression networks, which have enhanced power to identify co-regulated sets of genes (modules) that are likely to be involved in common biological processes. We constructed co-expression networks on the AD patients for each dataset after filtering out lowly expressed genes (Online Methods). The MAYO-microglial co-expression network consists of 45 modules ranging in size from 32 to 2,201 gene members. The ROSMAP-microglial co-expression network consists of 46 modules ranging in size from 115 to 892 gene members. In comparing all pairs of modules between the datasets for overlap, we identified 133 module pairs with significant overlap (FDR<0.05 for Fisher’s Exact Test, Figure 3A).

**Figure 3.**
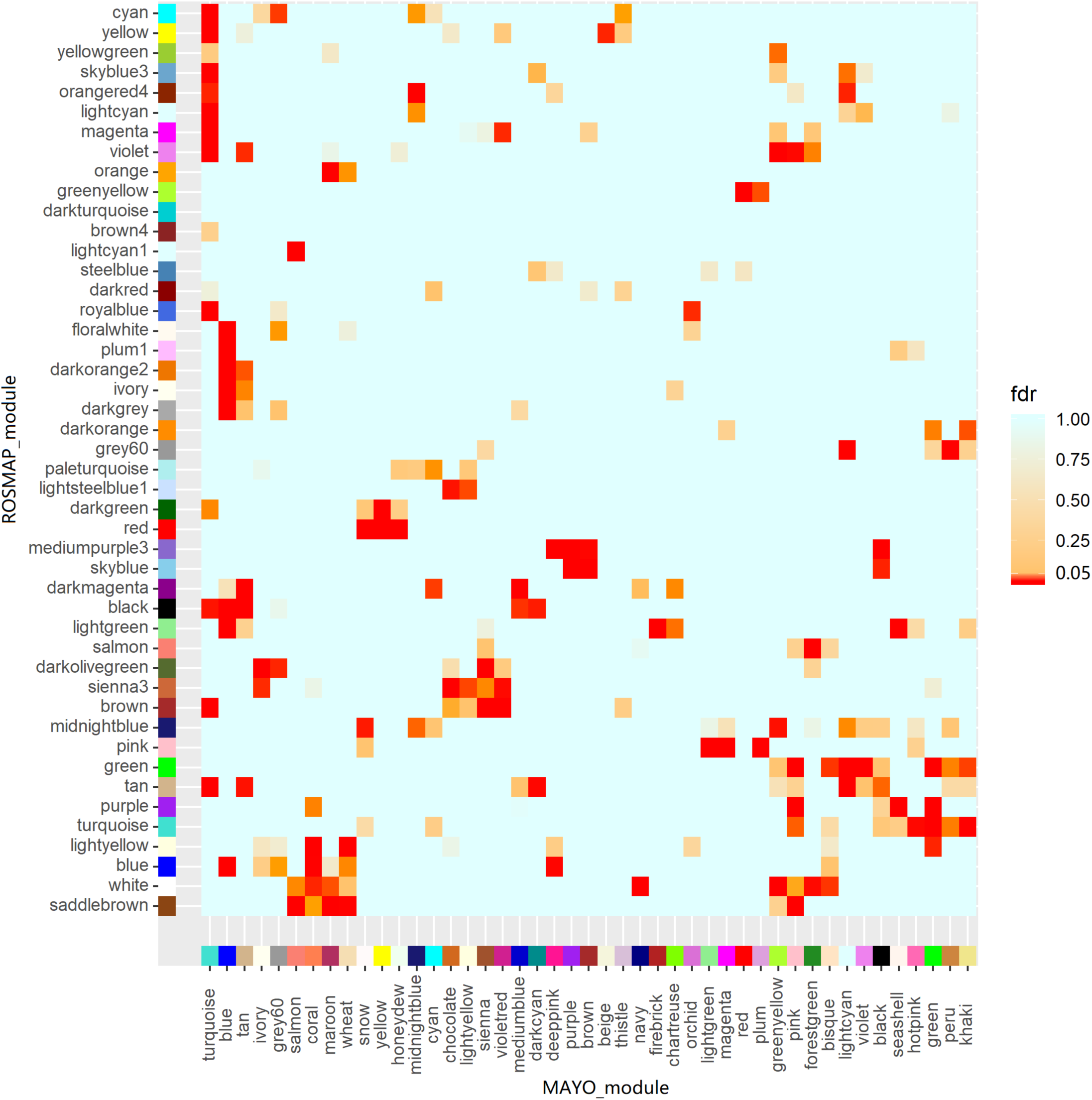
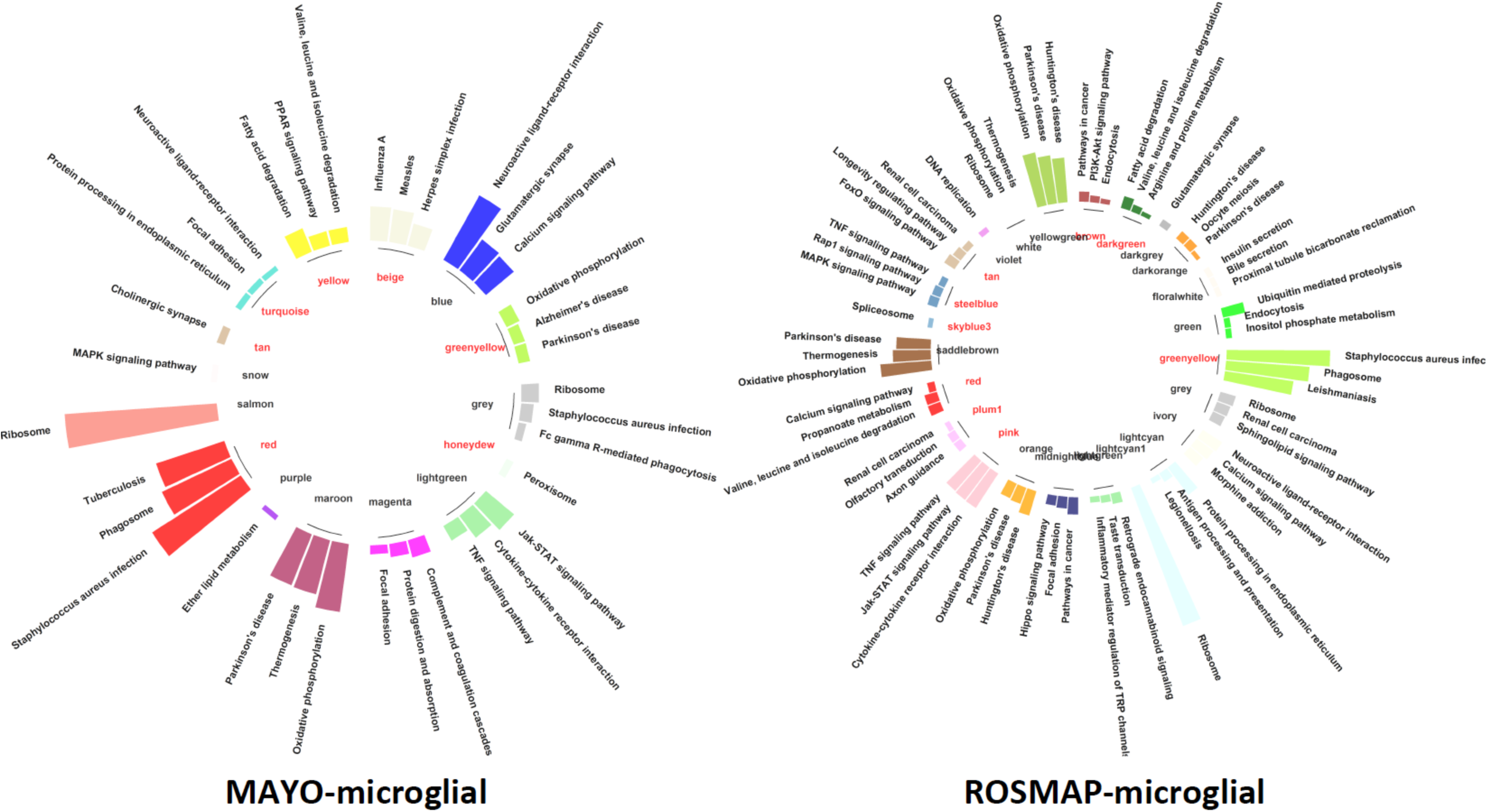

To characterize the functional relevance of the microglial-specific modules to AD pathology, two measures were employed: 1) Fold-enrichment for AD DE genes in each module; 2) Fold-enrichment for known microglial cell marker genes[31]. From these measures, we selected a seeding set of genes for input into the causal predictive network modeling by pooling genes from microglial-specific modules: 8 modules from the MAYO-microglial co-expression network (turquoise, pink, greenyellow, tan, mediumblue, darkcyan enriched for AD microglial DE genes; red and beige enriched for microglial cell markers) and 11 modules from the ROSMAP-microglial co-expression network (black, steelblue, brown, yellow, tan, plum1, darkgreen, skyblue3, red, sienna3, and mediumpurple3 enriched for AD microglial DE genes; pink and greenyellow enriched for microglial cell markers).

It is known that microglial cells interplay with other brain cells, such as astrocytes in modulating amyloid pathology in mouse models of Alzheimer’s disease[83]. We note that 2 modules (red and darkgreen) are enriched for both microglial-DE signature and astrocyte markers, 1 module (plum1) is enriched for both microglial-DE signature and neuron markers, and 2 modules (mediumpurple3, tan) are enriched for both microglial-DE signature and oligodendrocyte markers from the ROSMAP-microglial co-expression networks, indicating that these interactions were reflected in our microglial-specific data. Failing to account for these interactions will result in a compromised network model, however, over-counting these interactions will decrease the network specificity to microglial cells. Therefore, we only considered microglial-interacting cell types whose marker genes are the most significantly enriched (FDR-value<10E-4) by the same modules that are also the most significantly enriched (FDR-value<10E-4) for microglial-DE signature (Figure S7). Consequently, on top of the selected modules above, we added 2 modules (yellow and honeydrew) and 2 modules (darkgreen and red) enriched for astrocyte cell marker genes from MAYO-microglial and ROSMAP-microglial co-expression networks. Genes in these additional modules reflecting astrocyte-microglia interactions comprised only 13.9% (583 genes) and 16.9% (873 genes) of the MAYO- and ROSMAP-microglial networks, respectively.

To further annotate the co-expression modules selected above with respect to the biological processes they participate in, we performed pathway enrichment analysis to identify overrepresented biological processes across all selected modules in each dataset. Of the 10 and 13 selected modules from the MAYO- and ROSMAP-microglial networks, 7 and 9, respectively were significantly enriched (FDR<0.05) for KEGG pathways (highlighted red in Figure 3B). There are 104 and 142 significant pathways enriched by the selected modules from the MAYO and ROSMAP networks with an overlapping of 78 significant pathways (Fish’s Exact Test, OR=7.445, p-value=1.92E-15, Supplementary File S1), including the phagosome pathway.

### Predictive Network Modeling of Genetic Regulations Identified Pathological Pathways and Key Drivers for Microglial Function in AD

The ultimate goal of this study was to identify upstream master regulators (referred to here as key drivers) and pathways in microglia that contribute to AD pathology. Based on our DE, eQTL and co-expression network analysis, we built causal predictive network models on the subset of genes comprising co-expression network modules selected above by integrating the eQTLs and microglial-specific RNA-seq data. To this end, we developed a novel top-down & bottom-up predictive network modeling pipeline (Online Method) and applied it to the microglia-specific gene expression data in AD. The bottom-up component of our network reconstruction pipeline incorporates known pathway/network relationships derived from the literature and other sources, while the top-down component represents a structure-based learning algorithm that infers causal relationships supported by the eQTL and gene expression data.

To build the predictive network, we first pooled all genes from the subset of 10 and 13 selected modules in the MAYO- and ROSMAP-microglial co-expression networks respectively, to derive a set of seeding genes for each cohort (4187 for MAYO-microglial and 5152 for ROSMAP-microglial co-expression networks). The overlap between the two seeding gene sets contains 1842 genes (35.7% of ROSMAP and 43.9% of MAYO). Therefore, analysis using these two datasets increases the power to build robust networks and to discover high-confidence microglial key drivers that are associated with AD pathology. As *cis*-eQTLs causally affect the expression levels of neighboring genes, they can serve as a source of systematic perturbation to infer causal relationships among genes[51, 84, 85]. Consequently, we incorporated *cis*-eQTL genes into each network as structural priors. Of 5186 and 3875 unique *cis*-eQTL genes identified in the ROSMAP- and MAYO-specific datasets, 1978 and 687 overlapped with genes in each network respectively. The final causal predictive networks were comprised of 4600 and 4008 genes in the ROSMAP- and MAYO-microglial predictive networks (Figure 4A) respectively, with 1646 genes overlapping each network.

**Figure 4.**
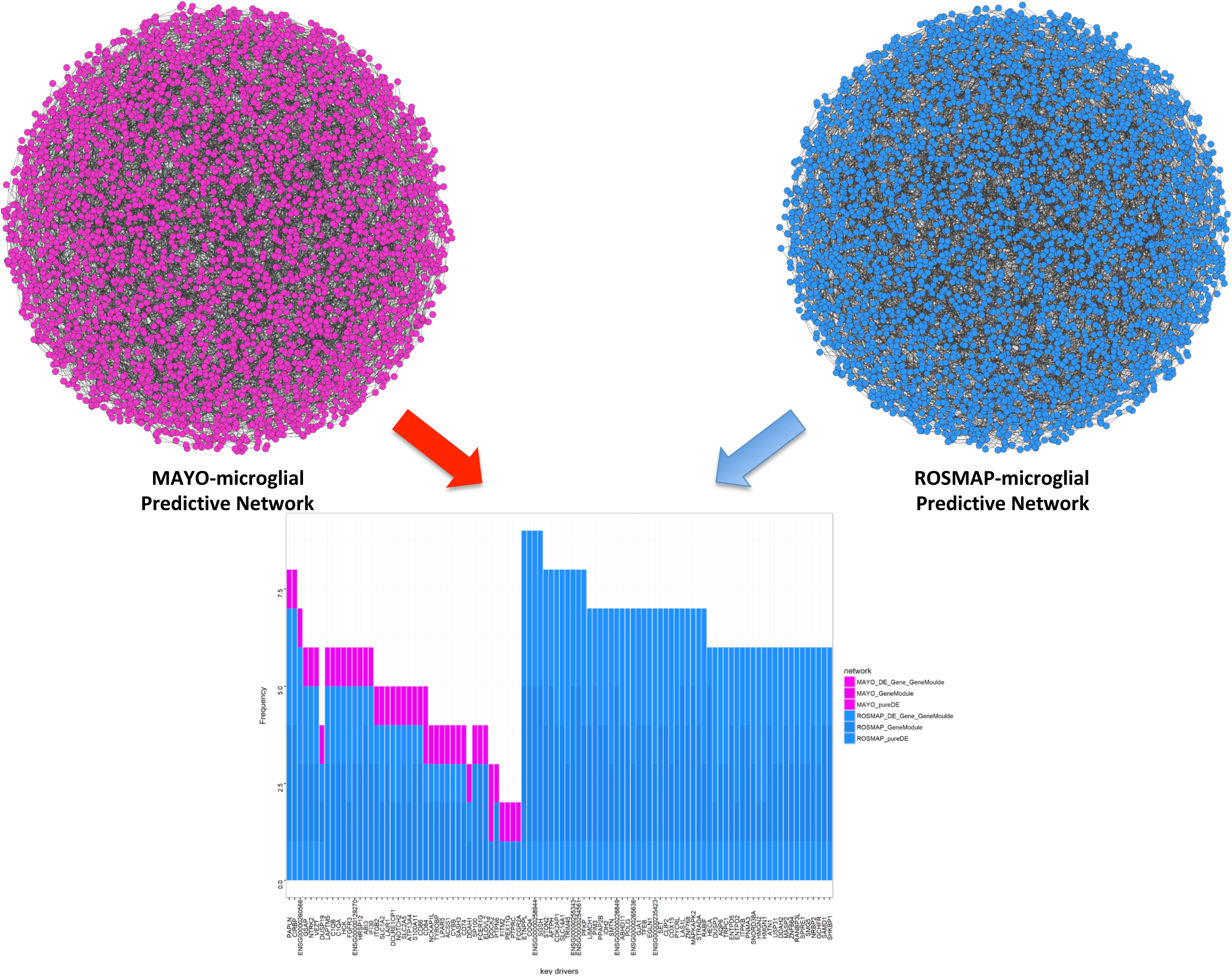
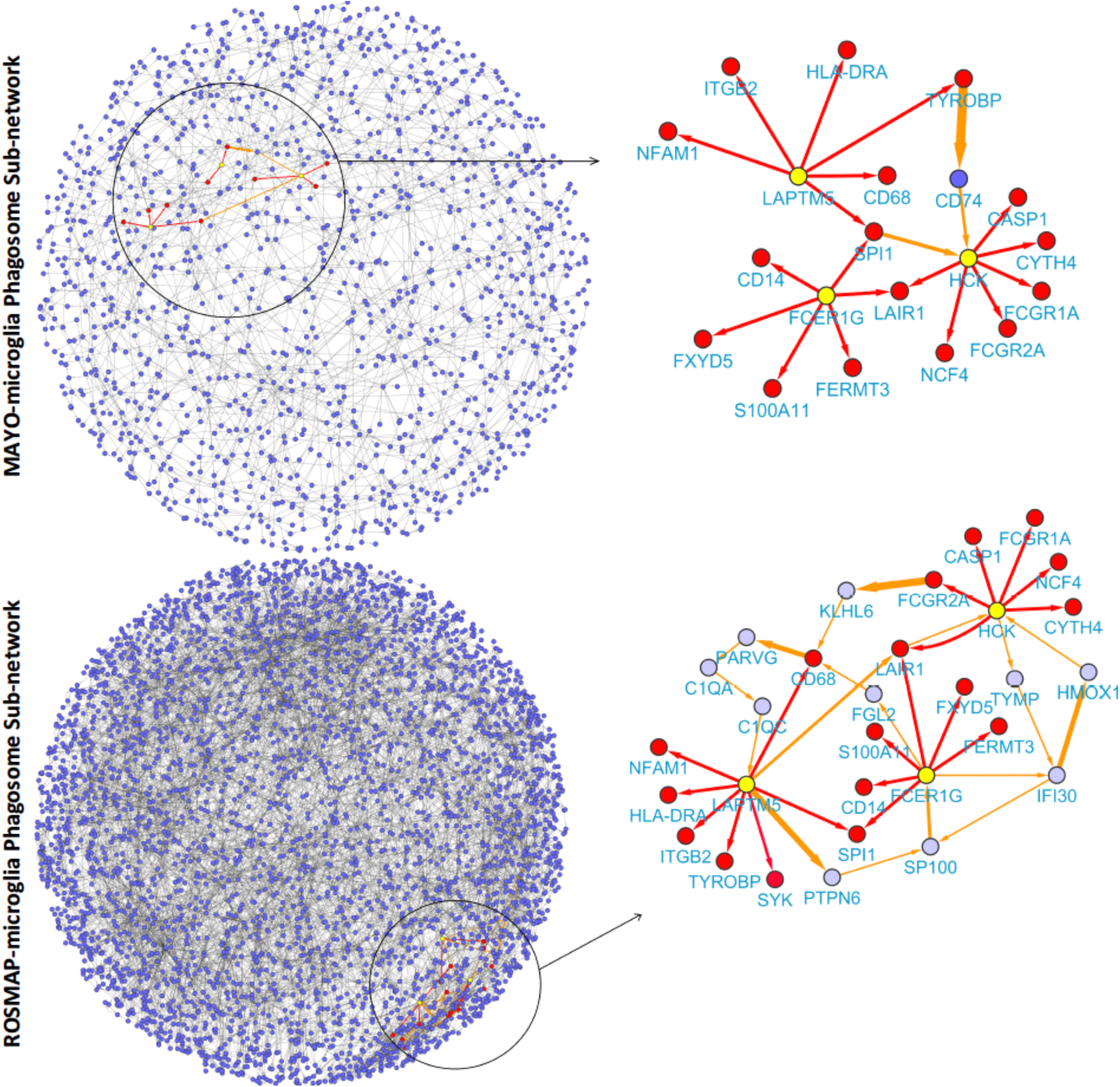
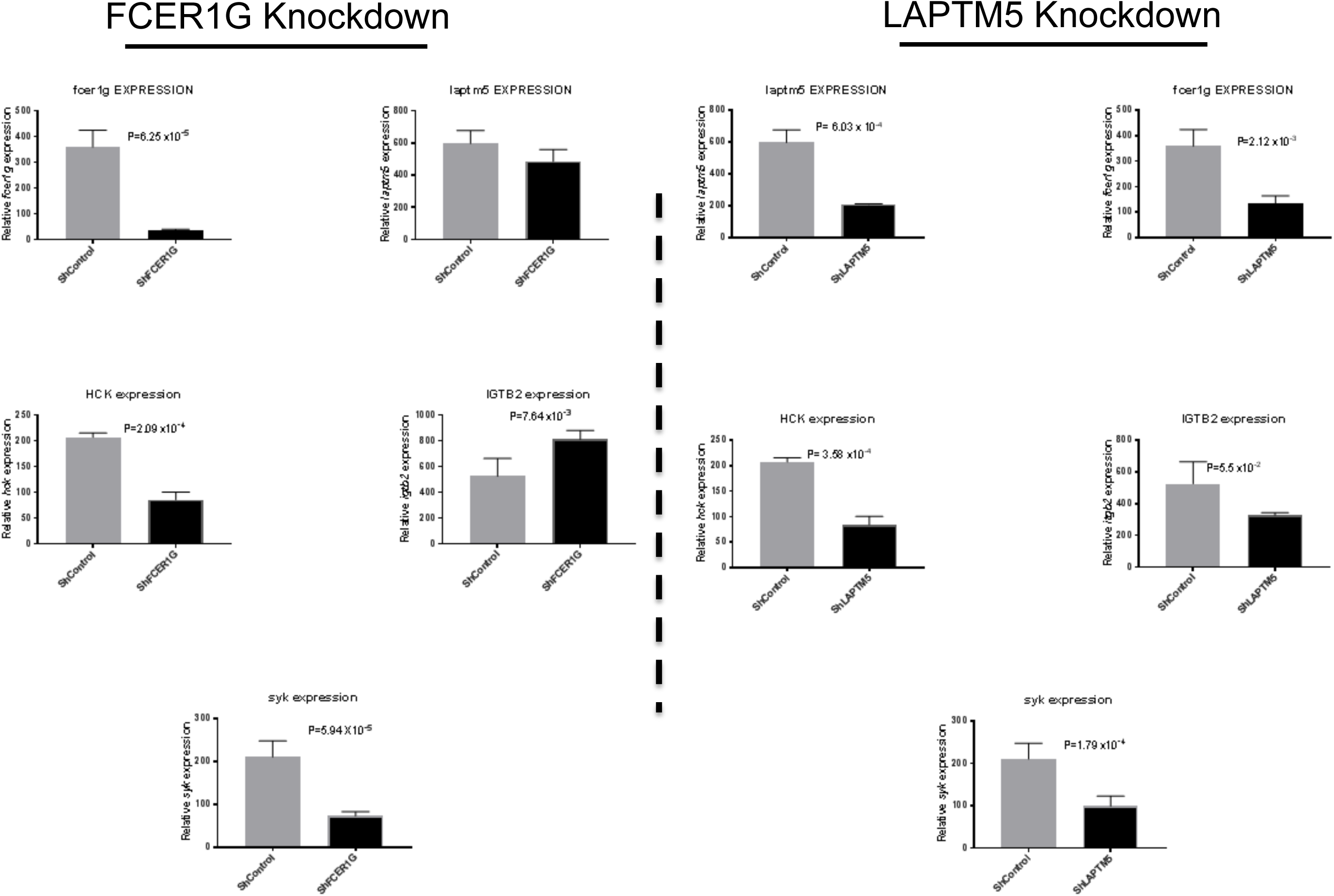

### Identification of Microglial-specific Key Drivers indicate the Phagosome Contributes to AD

Given the predictive networks, we applied Key Driver Analysis (KDA, Online Method) to derive the list of Key Driver (KD) genes for each network (Figure 4A). KDA seeks to identify those genes in the causal network that modulate network states. In the present analysis, we applied KDA to microglial-specific predictive network models to identify those genes causally modulating the states of these networks. In total, we identified 757 and 164 KD genes identified in the ROSMAP- and MAYO-microglial networks respectively and 43 KDs replicated across both networks. We prioritized all key drivers based on i) whether a gene predicted as KD is replicated across both datasets; ii) the number of different categories of target source predicting a gene as KD used by KDA (In this study, we used 3 kinds of targets for KDA: DE genes, modules and overlapping genes of DE gene with modules); and iii) the number of different target sources used by KDA predicts a gene as a KD.

Among the pathways enriched by the 43 KDs (Figure S8), *Microglial Pathogen Phagocytosis Pathway* is the most significantly enriched (p-value=2.60E-12) biological function by these 43 replicated KDs. We identified three KD genes, *FCER1G*, *HCK* and *LAPTM5* which are known to be involved in the phagosome as driving the AD phenotype, indicating that microglia-mediated phagocytosis causally links to the AD phenotype. Though these three genes are known to be involved in phagocytosis, their molecular mechanism in phagocytosis and the causal relationship among them is still unclear. To illustrate the molecular mechanism of these genes in phagocytosis, we extracted the downstream sub-network of these three KDs in the ROSMAP- and MAYO-microglial networks respectively (Figure 4B), and then performed CPDB pathway enrichment analysis[86], which confirmed that both sub-networks are significantly (p<0.05) enriched for Phagocytosis and Phagosome Pathways (Supplementary File S2). In addition, we predicted their causal relationship with the AD phenotype. In the MAYO-microglial sub-network, there are a total of 189 enriched pathways, including Microglia Pathogen Phagocytosis Pathway (p-value=0.0167), ER-Phagosome pathway (p-value=0.018) and Phagosome (p-value=0.028). In the ROSMAP-microglial sub-network, there are 534 enriched pathways, including Microglia Pathogen Phagocytosis Pathway (p-value=6.06E-11), Cross-presentation of particulate exogenous antigens (phagosomes) (p-value=1.81E-05), Fc gamma R-mediated phagocytosis (p-value=1.94E-05), Fc-gamma receptor (FCGR) dependent phagocytosis (p-value=4.03E-04), Role of phospholipids in phagocytosis (p-value=9.55E-04), Phagosome (p-value=2.24E-03), ER-Phagosome pathway (p-value=0.03). The overlapping contains 77 pathways, including Microglia Pathogen Phagocytosis Pathway, Phagosome, and ER-Phagosome pathway.

### Validation of the microglial key driver and network for AD pathology by knockdown in monocyte-derived microglia-like cells

To validate the molecular mechanism and pathways of these three key drivers captured by our predictive network model, we first sought to validate the downstream and upstream genes of microglial-specific key drivers (*HCK*, *FCER1G*, *LAPTM5*) predicted by our network model. To address this, we utilized the previously characterized highly efficient *in vitro* cell model system composed of human monocyte-derived microglia-like cells (MDMi) that recapitulates key aspects of microglia phenotype and function[33]. Using this model, we applied three different constructs of short hairpin lentiviral knockdown vectors targeting different parts of each KD gene by RNA interference (shHCK, shFCER1G and shLAPTM5) to MDMi cells differentiated from 3 healthy donors. We then picked two of the three constructs that gave at least 70-90% knockdown in the gene of interest compared to a control shRNA as verified by qPCR (Figure S9). To validate the predictive network, we measured the gene expression between MDMi cells that received the shHCK, shFCER1G or, shLAPTM5 and empty vector (shCTRL) for six network-predicted immediate downstream genes for *HCK*, *FCER1G* and *LAPTM5* respectively using Taqman real-time PCR (Figure 5A). Out of all immediate downstream genes of the KDs predicted by the two microglia networks, the six downstream genes were chosen on the basis of their expression level in the MDMi cells as determined by RNA-seq data set on MDMi (Supplementary File S3). For *HCK*, we selected *CASP1*, *FCGR1A*, *CYTH4 and NCF4*, *LAIR1*, *FCGR2A* from the MAYO- and ROSMAP-microglia networks respectively. Of the six genes, knockdown of the *HCK* gene in MDMi cells led to statistically significant reduction (p-value<0.05) (Online Method) in gene expression of five downstream genes in at least one construct: *CYTH4*, *NCF4*, *LAIR1*, *FCGR1A* and *FCGR2A* as compared to MDMi cells that received the control vector (ShCTRL). There was no significant decrease in gene expression from *CASP1* in either construct, though they trended in the same direction (p-values=5.87E-02 and 6.62E-02). For *FCER1G*, we selected *FERMT3*, *CD14*, *SPI-1*, *LAIR1* and *S100A11*, *FXYD5* from the MAYO- and ROSMAP-microglia networks. Five out of six downstream genes showed a significant reduction in gene expression in shFCER1G MDMi cells as compared to MDMi cells that received the control vector (ShCTRL) in at least one construct. *S100A11* showed no significant decrease in gene expression with p-values (5.68E-02 and 7.69E-02) close to the significance threshold. Similarly, we also conducted experiments using shRNA targeting the third key driver *LAPTM5*. For *LAPTM5*, we selected *NFAM, TYROBP, HLA-DRA, CD68 and NFAM, TYROBP, ITGB2, SPI-1* from MAYO- and ROSMAP-microglia networks. Knockdown of *LAPTM5* significantly upregulates *ITGB2*, *HLA-DRA* and *CD68*, and significantly downregulates *SPI-1*. Furthermore, knockdown of *LAPTM5* in MDMI cells did not have any significant effect on the expression of *TYROBP* (p-value=7.61E-01 and 5.44E-01) and *NFAM* (p-value=7.27E-02 and 6.07E-02), which is close to the significance threshold. *TYROBP* in our network contains 2 and 5 parent genes respectively in the MAYO- and ROSMAP-microglia networks. In the ROSMAP-microglia network, it has significantly more parents than all other nodes in the network (p-value=2.2e-16), potentially explaining why modulation of *LAPTM5* does not regulate *TYROBP* in our model system. In addition, we measured the gene expression between MDMi cells that received the shHCK, shFCER1G or, shLAPTM5 and empty vector (shCTRL) for four network-predicted common upstream genes (*C1ORF162*, *CKLF*, *SLC37A2*, *RMDN1*) for the three KDs respectively using Taqman (Figure 5B). As predicted by the network, none of these genes are significantly changed. The experimental results are listed in Supplementary File S4 (sheet ‘KD, downstream genes, results’).

**Figure 5.**
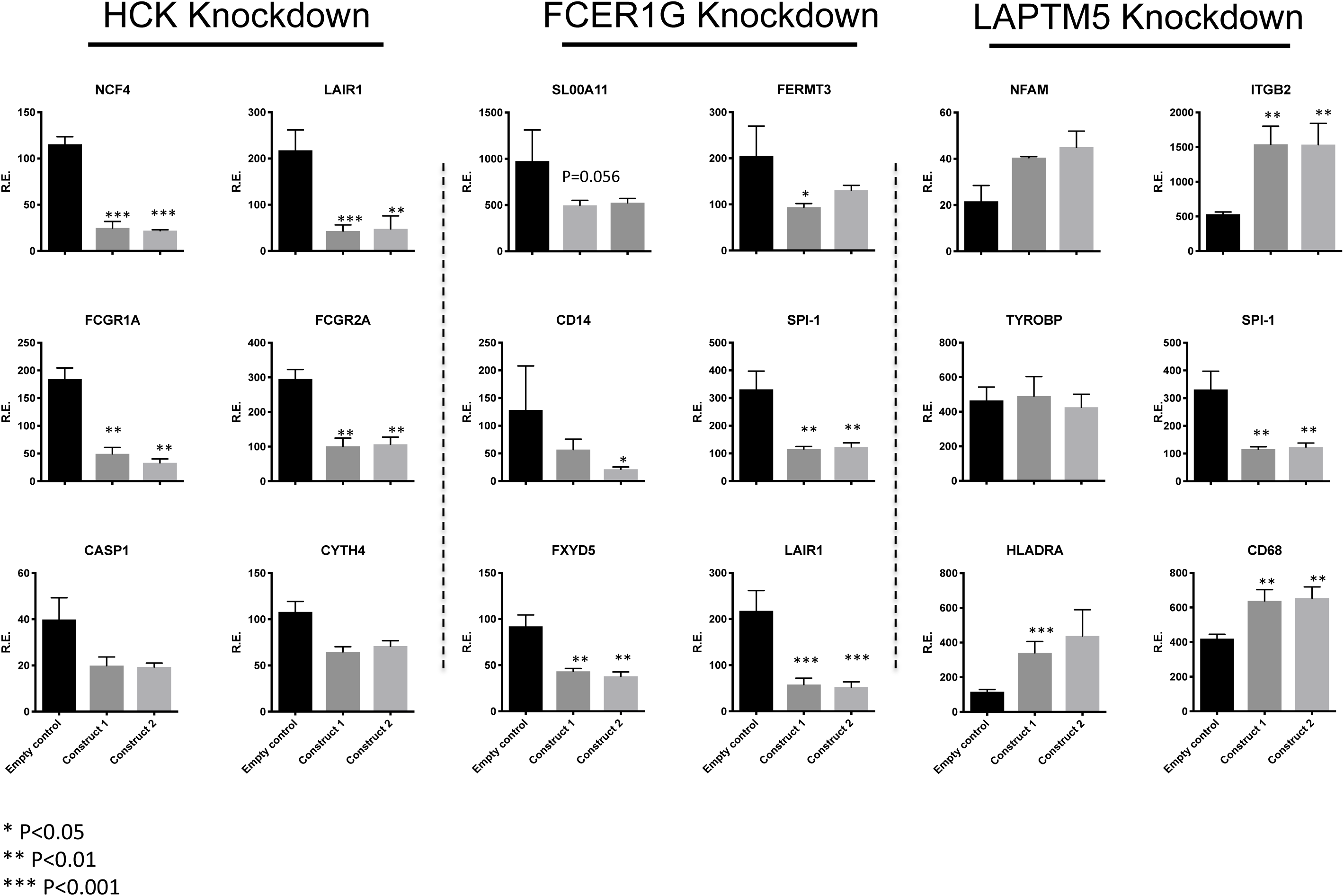
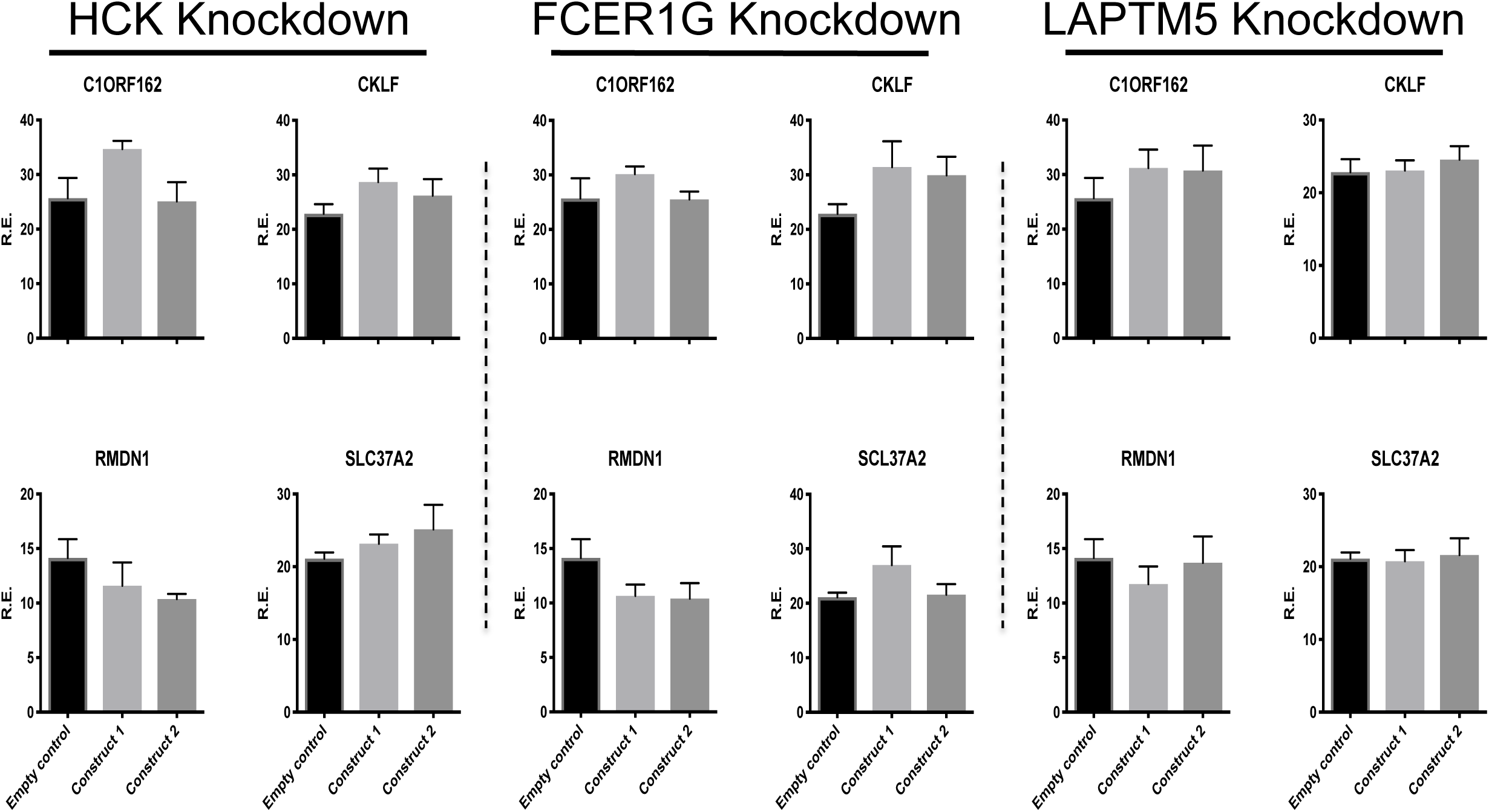
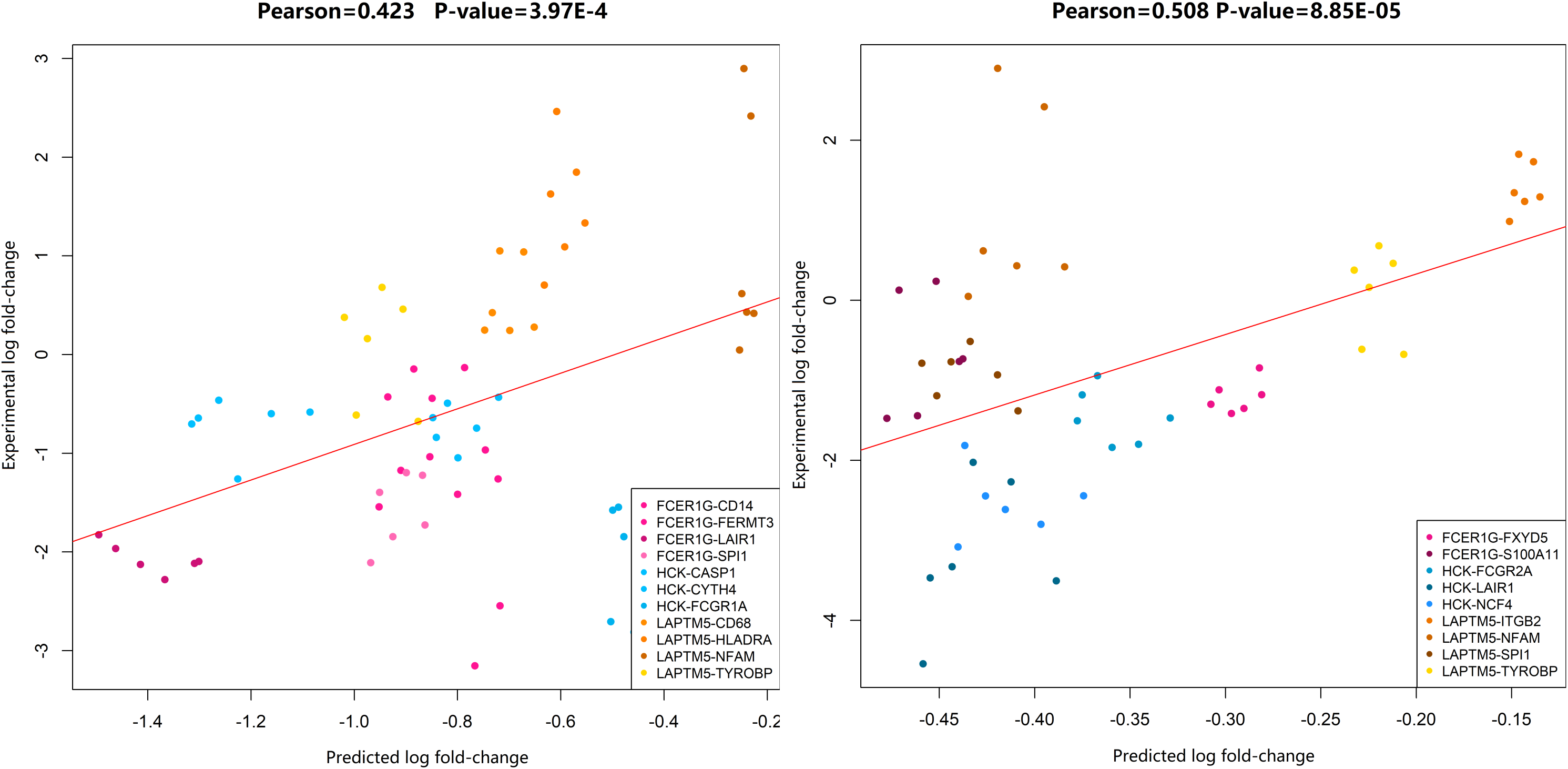
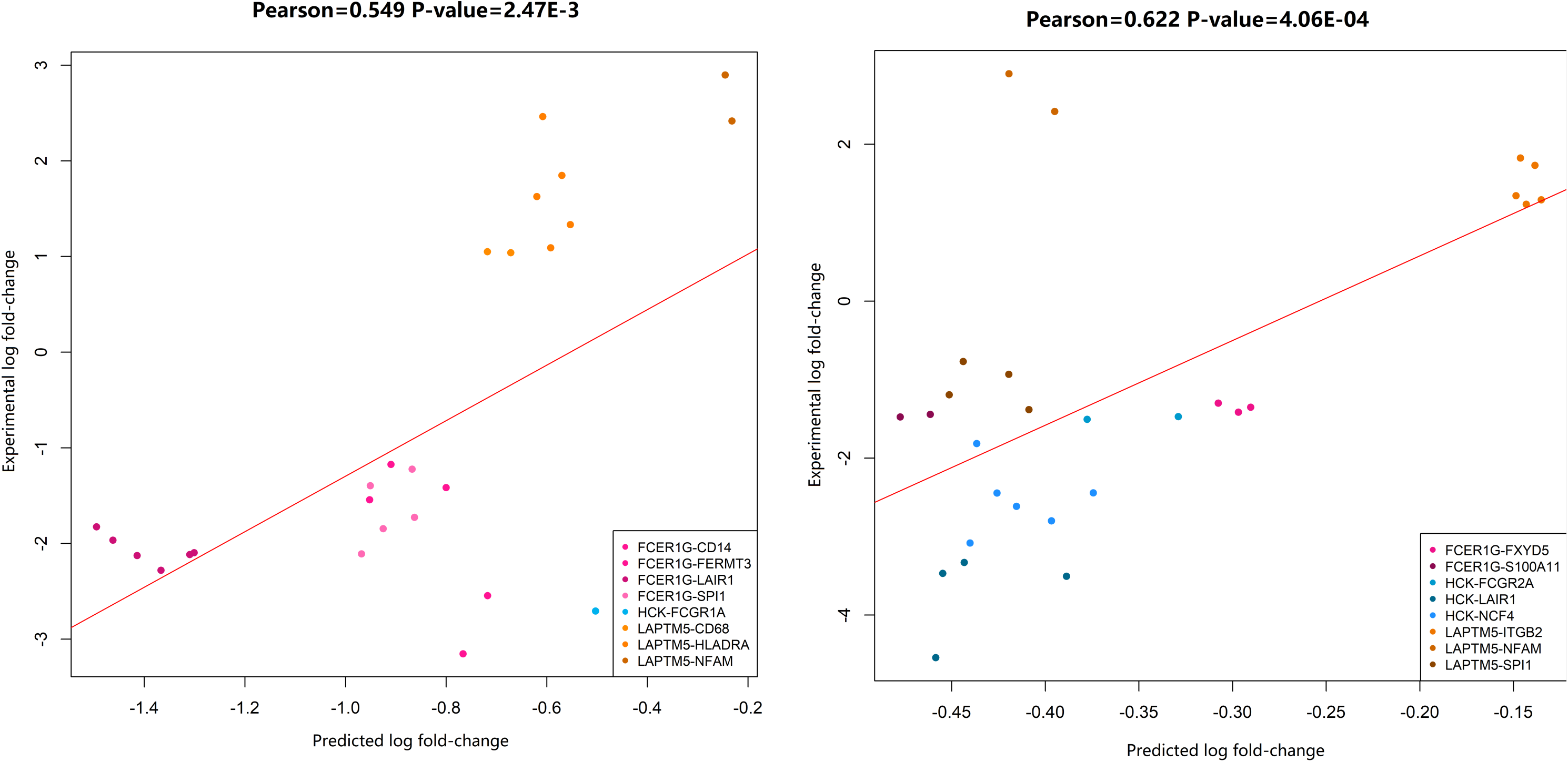

### Validation of In-silico Prediction on Gene Expression Prediction by Microglial-specific Predictive Networks

Next, we predicted in-silico gene expression of downstream genomic targets from these three key drivers based on our predictive network model. We developed the in-silico phenotypic prediction pipeline, which is a component of the predictive network, to predict i)qualitative response, i.e. direction of gene expression fold-change and ii)quantitative response, i.e. expression fold-change of the downstream genes of three key drivers in the ROSMAP and MAYO-microglial networks. In shRNA experiments, we measured gene expression fold-change of 11 and 9 KD-target pairs (see last section) predicted by the MAYO- and ROSMAP-microglial networks with two constructs in MDMi cells from 3 healthy donors. Therefore, there are a total 66 and 54 measurements in the MAYO- and ROSMAP-microglia network respectively. First, we predicted the direction of gene expression change (up-regulation or down-regulation) and compared it to those in the shRNA experiments. Out of the 66 measurements, all predicted to be down-regulated by the MAYO network, 22 measurements were up-regulated and 44 were down-regulated in the shRNA experiment. Out of 54 measurements predicted to be down-regulated by the ROSMAP network, 18 were up-regulated and 36 were down-regulated in the shRNA experiment. The accuracy of the qualitative response prediction is 66.7% for both networks. Next, we calculated the Pearson correlation (0.423 and 0.508, p-value=3.97e-04 and 8.85e-05) and Spearman correlation (0.367 and 0.436, p-value=2.61e-03 and 1.1e-03) between predicted and experimental expression fold-change (Online method, Supplementary File S5) in the MAYO- and ROSMAP-microglia networks (Figure 5C). Further, if we only consider measurements (p-value<0.05) in shRNA experiments, 28 and 28 pairs are used to calculate the Pearson correlation (0.549 and 0.622, p-value=2.47e-03 and 4.06e-04) and Spearman correlation (0.358 and 0.492, p-value=6.21e-02 and 8.51e-03) in the MAYO- and ROSMAP-microglia networks (Figure 5D). This result demonstrated that our predictive network model accurately predicts downstream gene expression changes upon perturbing any key driver, which is validated using Taqman.

### Validation of the causal regulation among HCK, FCER1G and LAPTM5 by knockdown in MDMi cells

Next, we identified previously unknown causal regulation among the three key drivers in the process of phagocytosis using our microglia-specific predictive network models in ROSMAP and MAYO. We extracted sub-networks of phagocytosis around the 3 key drivers (highlighted in Figure 4B), and determined that *HCK* was downstream of *FCER1G*, which is, in turn, downstream of *LAPTM5*. These causal regulations are replicated in MAYO- and ROSMAP-microglial networks. Further, to validate the relationship between HCK, FCER1G and LAPTM5, we applied shRNA targeting FCER1G and *LAPTM5* using lentivirus in MDMi cells. We then measured *HCK*, *LAPTM5* and *FCER1G* expression in each knockdown. Gene expression data from the shFCER1G cells showed significant reduction (p-value=2.09E-4) in *HCK* gene but there was no significant (p-value=1.62E-01) change in LAPTM5 expression. In addition, gene expression data from shLAPTM5 showed significant reduction in HCK (p-value=3.58E-04) and FCER1G (p-value=2.12E-03) (Figure 4C), thereby indicating that LAPTM5 was upstream of FCER1G and HCK was downstream of FCER1G, which validated our microglia-specific network. Besides HCK and FCER1G, ITGB2 and SYK gene are also significantly changed their expression level in both knockdown experiments, suggesting that they are directly or indirectly downstream of LAPTM5 and FCER1G (Figure 4C). Interestingly, both our MAYO and ROSMAP sub-networks show that ITGB2 is downstream of LAPTM5, and in the ROSMAP sub-network, ITGB2 is an indirect downstream gene of FCER1G. In addition, in the ROSMAP sub-network, SYK is direct downstream gene of LAPTM5 and indirect downstream gene of FCER1G (Figure 4B). The experimental results are listed in Supplementary Table S4 (sheet ‘KD, subnetworks, results’).

### Functional Validation of HCK and FCER1G as key drivers of microglia function of Aβ clearance

Microglia are innate immune cells of the brain which play an important role in clearance of amyloid-beta which forms plaques in the brain, a hallmark pathology of AD. In this study, we identified key drivers in our predicted microglia-specific causal network model for phagocytosis (HCK and FCER1G) and lysosome function (LAPTM5). In order to validate the function of HCK and FCER1G as mediators of phagocytosis, we analyzed the Aß42 uptake ability of MDMi cells differentiated from 4-5 healthy donors that received lentivirus containing the short hair pin RNA targeting HCK (shHCK) and FCER1G (shFCER1G) and compared it to the MDMi cells from the same donors that received empty control virus (shCTRL) using fluorescently labelled Aß42 peptide. The MDMi cells with both shHCK and shFCER1G show a significant decrease in Aß42 uptake for the two *HCK* constructs (paired t-test, p-value = 0.030 and 0.028) and for the two *FCER1G* constructs (p-value=0.037 and 0.045 (Figure 6). As *LAPTM5* is a lysosomal molecule, we do not expect to see an effect in our uptake assay. Thus, we functionally validated these genes as key drivers of phagocytosis in microglia cells as predicted by our network specific model. The complete results are in Supplementary File S4 (sheet ‘Abeta uptake, results’).

**Figure 6.**
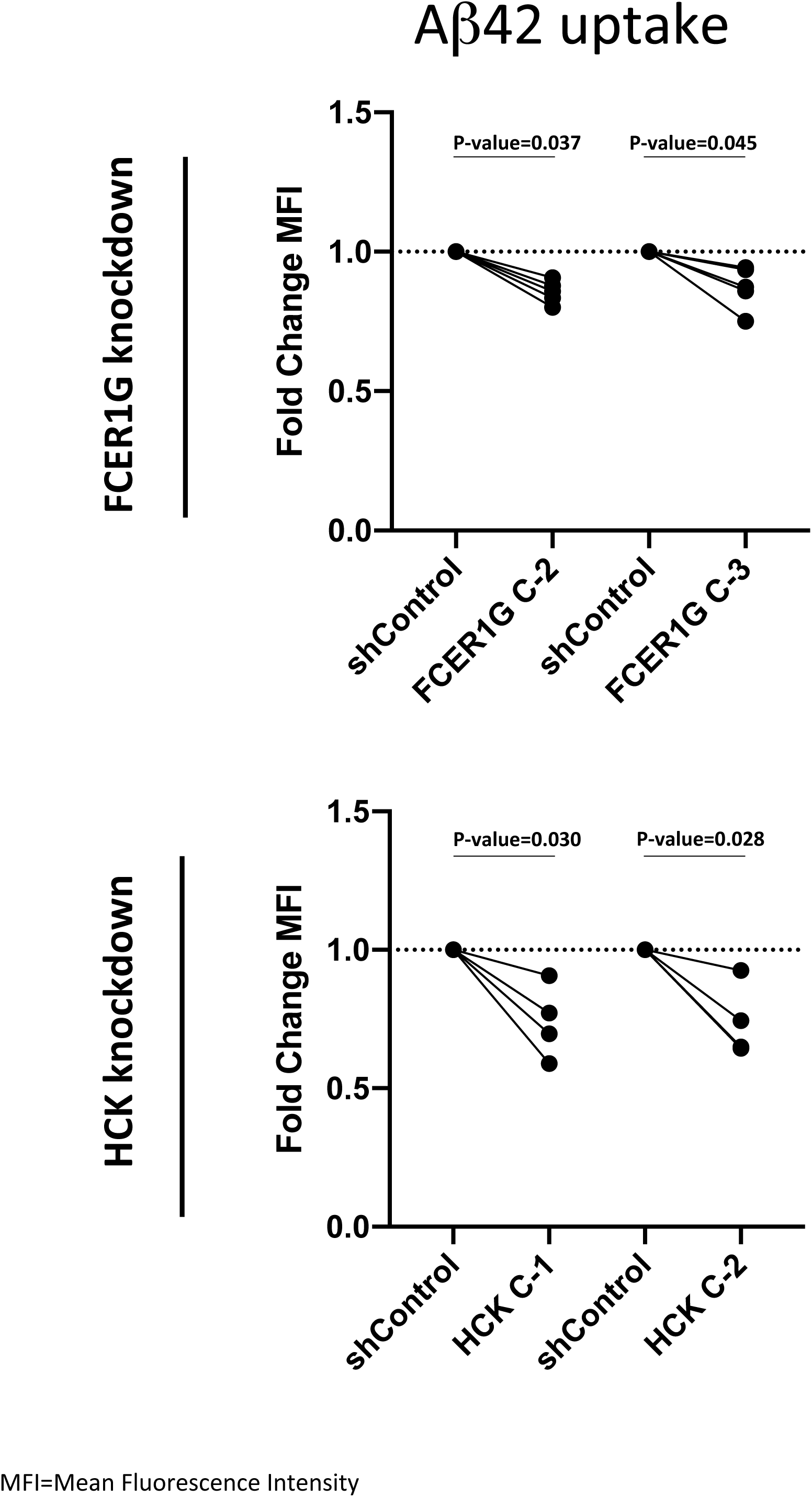

## Discussion

GWAS studies in LOAD have identified several microglial specific genes. However, it is unclear how these genes interact and what cellular pathways are involved in the pathology of LOAD. Hence, a comprehensive characterization of gene regulatory networks with association to disease is important to provide insights into the underlying mechanisms of complex neurodegenerative diseases such as Alzheimer’s disease. Our study uses an innovative predictive computational systems biology model to identify upstream regulators (key drivers) and cellular pathways in microglial cells that contribute to AD pathology using the Mayo Clinic and ROSMAP datasets in the AMP-AD consortium. The RNA-seq data from brain-region tissue in MAYO (TCX-Temporal Cortex, 79 AD and 76 CN) and ROSMAP (PFC-Pre-frontal Cortex, 212 AD and 200 CN) cohorts are computationally de-convoluted into single cell types including neurons, microglia, astrocytes, endothelial cells and oligodendrocytes. In this study, we focused on microglial-specific gene expression data. We performed preliminary bioinformatics analysis including differential expression, eQTLs, co-expression networks and pathway analysis prior to building predictive causal network models. There are 4187(43.9%)/5152(35.7%) (overlap percentage in parenthesis) genes in the seeding gene list of MAYO-/ROSMAP-microglial predictive network models and 4008(41.0%)/4600(35.7%) genes in the final MAYO-/ROSMAP-microglial predictive network models. The difference of key drivers/pathways in each predictive network can be attributed to: i) different susceptibility and pathology response to AD in TCX and DLPFC regions analyzed in the MAYO and ROSMAP cohorts respectively; ii) different patient composition in the two cohorts, such as Male/Female, APOE4+/- and clinical stage; and iii) technical variance in sample extraction, RNA preparation, and RNA-sequencing as well as other covariates. However, the replicated key drivers/pathways derived by these network models identified robust biological processes and key drivers in microglial cells under AD diagnosis despite the significant variance in data and cohorts as described above. Consequently, our predictive networks identified robust key drivers for phagocytosis in microglial-specific cells associated with AD, which are validated using an *in vitro* model of microglia.

Our novel predictive network-based analysis integrated microglial cell-specific genetics and genomics data to identify key regulatory genes associated with microglial functions in AD, i.e. *LAPTM5*, *FCER1G* and *HCK*. Pathway enrichment analysis confirmed that all three key drivers and their downstream genes in the network model regulate phagosome processes. The phagosome pathway is activated upon neuronal loss or amyloid plaque buildup, two important pathophysiological hallmarks of AD. Microglia clear synapses and neurites during development and in neurodegenerative processes using phagocytosis via C1Q, C3, CR3, and the DAP12/TYROBP cascade[87–89]. Microglia are also closely associated with amyloid beta plaques making it important to understand the cause-and-effect relationship between immune cells and AD progression[90].

Our network model predicted regulation between FCER1G, HCK and LAPTM5 genes. We highlight that HCK is downstream of FCER1G in the network with our lentiviral shRNA knockdown in MDMi cells (Figure 4C). A study by Taguchi et al[36] showed that HCK and FCER1G are up-regulated in the cortex of *App*^NL-G-F/NL-G-F^ mice as Aβ amyloidosis progressed thereby associating them with plaques and phagocytosis. Furthermore, FCER1G shows significant association in AD, in regards to immune and microglial functions and amyloid deposits in humans and mice[34–36]. The reduction in uptake of the fluorescently labeled amyloid beta 1-42 in shFCER1G and shHCK microglia cells indicates the significance of these genes as phagocytic modulators in microglia cells, which validated the accuracy of our predictive network model. Furthermore, while *LAPTM5* is upstream of *FCER1G* and *HCK*, its main role is in lysosomal function, which is not captured in our assay, and thus modulating *LAPTM5* doesn’t show a robust effect on uptake of Aß42 (Supplementary File S4, sheet ‘Abeta uptake, results’).

GWAS studies have implicated SNPs and polymorphisms in the TYROBP binding protein TREM2, ITGAM and SPI-1 genes as being associated with LOAD[91]. Using our computational predictive network, we also highlight that TYROBP is downstream of LAPTM5 and TREM2 and hence is regulated by both genes, while ITGAM is upstream of LAPTM5, FCER1G and HCK. We further demonstrated that SPI-1, an established GWAS loci for AD, is a downstream gene of our key driver LAPTM5 in a microglial specific network from the ROSMAP/MAYO data sets. Our study demonstrates the importance of these classical immune genes in AD functions.

Comparative profiling of human cortical gene expression in AD patients[35] and mouse models with amyloid beta plaque accumulation have shown involvement of our key drivers LAPTM5, FCER1G and HCK. These three genes were identified in Zhang et al. (ref.), however here we demonstrate that they exist in one causal network, and we have delineated the upstream and downstream relationships. A recent study by Lim et al. shows that inhibition of HCK dysregulates microglial function of phagocytosis and enhances amyloid plaque build-up in the J-20 mouse model of AD[92]. In addition, a recent study[41] showed that LAPTM5 is significantly associated with the mouse amyloid response network and that it’s human ortholog contains SNPs associated with AD.

In addition to network reconstruction and key driver discovery, our predictive network model is capable to perform in-silico phenotype prediction. Upon perturbing any number of genes in the network, we can predict i) whether a given gene in the network will significantly change their expression level; ii) the qualitative response, i.e. direction of gene expression change of the downstream genes to the perturbation; iii) the quantitative response, i.e. log fold-change of gene expression in the downstream genes to perturbation. We used two shRNAs targeting different regions of the three key driver genes and tested gene expression change of 18 downstream genes of these three key drivers in the network using Taqman array. Fourteen out of 18 (78%) predicted-to-change downstream genes by our network models are validated as significantly (p<0.05) altered in their gene expression by knockdown experiments. In addition, 4 common upstream genes of these key drivers predicted not-to-change by the networks, are 100% validated as not altering in expression after knockdown. The accuracy of the qualitative response prediction is 66.7% for the 18 downstream target genes in MAYO and ROSMAP-microglial networks. The Pearson correlation between experimental data and quantitative prediction by the model is very significant for all measured downstream targets in MAYO- and ROSMAP-microglial network respectively. This result demonstrated that our predictive network model is capable of further predicting phenotypic changes upon perturbations in the model.

Overall, our innovative computational systems biology modeling of microglia specific networks further deepens our understanding of microglial-specific implications in AD pathology by identifying robust causal networks of key driver genes and their genomic target genes for the phagosome, including a GWAS AD-related gene (LAPTM5) that has major implications in AD. Our predictive network not only identifies FCER1G, HCK and LAPTM5 as key drivers for microglial specific genes in the phagosome pathway but also demonstrates the functional association of HCK and FCER1G in amyloid beta uptake in microglial cells. Our approach appears to offer novel insights for drug discovery programs that can affect neurodegenerative diseases, such as LOAD.

## Author contribution

Conceptualization: R.C. E.M.B and E.E.S; ROSMAP RNA-seq and WGS pre-processing: N.D.M. M.Y.R.H.; MAYO RNA-seq and WGS pre-processing: M.A. X.W. J.S.R M. M.Y.R.H; Single Cell-type Gene Expression: R.C., M.Y.R.H., N.E.T; Data Analysis: R.C., M.Y.R.H., K.Z. S.M. M.L.A; MDMi shRNA experiment design and implementation: K.P., E.M.B. P.N. provided technical assistance. G.C., T.P., K.P. and E.M.B created and optimized the shRNA protocol. T.P. provided plasmids; Manuscript writing and figures: R.C. K.P. M.Y.R.C. K.Z. N.D.M; Manuscript Editing: E.M.B P.L.D. E.E.S. N.E.T. D.A.B.

## Acknowledgement

The authors thank the patients and their families for their participation, without whom these studies would not have been possible. This work was supported by the National Institute of Health: NIA R01 AG057457, NIA R01 AG059093 and NIA 1R01AG057931 to R.C; U01 AG046152, R01 AG048015, RF1 AG015819, R01AG056284, R01AG036836 R01AG043617 to PLD; NINDS R01 NS080820, NIA RF AG051504 and NIA U01 AG046139 to N.E.T; R01AG058852, R01NS089674, RF1AG057457, R01AG043617 to E.M.B; NIA U01AG058635 and NIMH R01MH109897 to E.E.S.

## Online Methods

### Data source

Data has been downloaded from the AMP-AD consortium database hosted on the Synapse.org data portal (doi:10.7303/syn2580853).

#### Mayo Clinic Transcriptome and Genome-Wide Genotype Data

The Mayo Clinic (MAYO) transcriptome and genome-wide genotype datasets utilized in this study have previously been described[1–4]. The Mayo Clinic temporal cortex RNA sequence data (Synapse ID: syn3163039) and genome-wide genotypes (Synapse ID: syn8650953) are available from AMP-AD Knowledge portal. Below, we provide details on these datasets:

#### Mayo Clinic Cohort Participants

Mayo Clinic RNAseq cohort has RNAseq-based whole transcriptome data from 278 TCX samples from subjects with the following diagnoses: 84 AD, 84 PSP, 80 controls and 30 pathologic aging. For this study, we utilized data from subjects with AD and controls. Subjects with AD had definite neuropathologic diagnosis according to the NINCDS-ADRDA criteria[5] and had Braak neurofibrillary tangle (NFT) stage of ≥4.0.

Control subjects each had Braak[6] NFT stage of 3.0 or less, CERAD[7] neuritic and cortical plaque densities of 0 (none) or 1 (sparse) and lacked any of the following pathologic diagnoses: AD, Parkinson’s disease (PD), DLB, VaD, PSP, motor neuron disease (MND), CBD, Pick’s disease (PiD), Huntington’s disease (HD), FTLD, hippocampal sclerosis (HipScl) or dementia lacking distinctive histology (DLDH). Within the Mayo RNAseq cohort, all AD and PSP subjects were from the Mayo Clinic Brain Bank. Thirty-one control TCX samples were from the Mayo Clinic Brain Bank, and the remaining control tissue was from the Banner Sun Health Institute. All subjects were North American Caucasians. All but control subjects, had ages at death ≥60, and a more relaxed lower age cutoff of ≥50 was applied for normal controls to achieve sample sizes similar to that of AD and PSP subjects. Brain samples for the Mayo RNAseq study underwent RNA extractions via the Trizol/chloroform/ethanol method, followed by DNase and Cleanup of RNA using Qiagen RNeasy Mini Kit and Qiagen RNase -Free DNase Set. The quantity and quality of all RNA samples were determined by the Agilent 2100 Bioanalyzer using the Agilent RNA 6000 Nano Chip. Samples had to have an RIN ≥5.0 for inclusion in the study.

All of this work was approved by the Mayo Clinic Institutional Review Board. All human subjects or their next of kin provided informed consent.

#### Mayo Clinic RNAseq Data

Mayo Clinic RNAseq samples were randomized across flowcells, taking into account age at death, sex, RIN, Braak stage and diagnosis. Library preparation and sequencing of the samples were conducted at the Mayo Clinic Medical Genome Facility Gene Expression and Sequencing Cores, as previously described[8]. The TruSeq RNA Sample Prep Kit (Illumina, San Diego, CA) was used for library preparation from all samples. The library concentration and size distribution was determined on an Agilent Bioanalyzer DNA 1000 chip. Three samples were run per flowcell lane using barcoding. All samples underwent 101 base-pair (bp), paired-end sequencing on Illumina HiSeq2000 instruments. Base-calling was performed using Illumina’s RTA 1.17.21.3. FASTQ sequence reads were aligned to the human reference genome using TopHat 2.0.12 [9] and Bowtie 1.1.0[10], and Subread 1.4.4 was used for gene counting[11]. FastQC was used for quality control (QC) of raw sequence reads, and RSeQC was used for QC of mapped reads. Raw read counts were log2-transformed and normalized using Conditional Quantile Normalization (CQN) via the Bioconductor package; accounting for sequencing depth, gene length, and GC content[12].

#### Mayo Clinic Genome-Wide Genotype Data

Subjects in the Mayo Clinic RNAseq cohort underwent whole genome genotyping using the Illumina Infinium HumanOmni2.5-8 BeadChip, which delivers comprehensive coverage of both common and rare SNP content from the 1000 Genomes Project (minor allele frequency>2.5%) providing genotypes for 2,338,671 markers. The genotyping was done at the Mayo Clinic Medical Genome Facility. Whole genome genotype calls were made using the auto-calling algorithm in Illumina’s BeadStudio 2.0 software, subsequent to which they were converted into PLINK formats for analysis[13].

#### Mayo Clinic RNAseq Data Quality Control (QC)

All Mayo Clinic TCX RNAseq samples had percent mapped reads ≥ 85%. Using R statistical software, raw read counts were transformed to counts per million (CPM), which were log2 normalized. Mean expression for chromosome Y genes with non-zero counts were plotted to identify any samples with deviation from expected expression based on recorded sex. Two AD TCX samples with discordant sex were excluded. Raw read counts were then normalized using Conditional Quantile Normalization (CQN) via the Bioconductor package; accounting for sequencing depth, gene length, and GC content. GC content was calculated via the Bioconductor package, Repitools and sequencing depth was calculated as the sum of reads mapped to genes. Genes with non-zero counts across all samples were retained and principal components analysis was performed using the prcomp function implemented using R Statistical Software (R Foundation for Statistical Computing, version 3.2.3). Principal components 1 and 2 were plotted and no outliers (>6SD from mean) were identified.

#### Mayo Clinic Genotype Data Quality Control (QC)

Genome-wide genotypes were obtained for all subjects in the Mayo Clinic RNAseq study using Illumina Omni 2.5 Beadchips. Samples were checked for discordant sex. The same two subjects that were excluded due to discordant sex based on RNAseq data were also determined to have discordant sex based on the genome-wide genotype data. Subjects were assessed for heterozygosity rate > 3SD from the mean. One AD sample with TCX RNAseq had high heterozygosity indicating possible sample contamination and 3 TCX RNAseq samples had low heterozygosity (2 controls and 1 AD) indicating either divergent ancestry or consanguinity. These samples were also excluded from the analysis. The dataset was filtered to include only autosomal SNPs. PLINK[13] --genome function was used to identify any sample duplicates or related pairs of subjects. Two pairs of samples were identified as > 3^rd^ degree relatives. For each pair, the sample with the lowest SNP call rate was excluded (1 PSP and 1 control). The dataset was further filtered to remove complex genomic regions (chr8:1-12,700,000; chr2:129,900,001-136,800,000; chr17:40,900,001-44,900,000; chr6:32,100,001-33,500,000) and LD pruned using the SNPRelate (v1.4.2) package in R (v3.2.3) [14], implementing an LD threshold of 0.15 and a sliding window of 1E-07 bp. Remaining SNPs and subjects were analyzed using EIGENSOFT[15] for population outliers. Two samples were identified as population outliers (1 PSP and 1 control) using the default parameters of > 6 SD from the mean on any of the top ten inferred axes following 5 iterations and were removed from further analysis.

Clinical, genotype and processed RNA-seq read count data was obtained privately from Dr. Nilufer Tan Taner lab at the Mayo Clinic.

#### ROSMAP Transcriptome and Genome-Wide Genotype Data

The ROSMAP dataset’s dorsolateral prefrontal cortex gene expression (RNA-seq BAM files), genotypes and clinical covariates were downloaded from synapse (respective synapse project IDs: syn4164376, syn3157325 and syn3191087) using the synapseClient R library (Matthew Furia (2015). synapseClient: Synapse R Client from Sage Bionetworks. R package version 1.11-1. http://www.sagebase.org). Requests for ROSMAP data can be made at www.radc.rush.edu.

#### ROSMAP Cohort Participants

*ROSMAP* dataset contains two cohorts: The Religious Orders Study (ROS) and The Memory and Aging Project (MAP)[16]. Both ROS and MAP are a longitudinal clinical-pathologic cohort studies of aging and dementia run by the Rush Alzheimer’s Disease Center. In both studies, participants enroll without known dementia and agree to annual clinical evaluation. All subjects agree to brain donation as a condition of entry. ROS enrolled individuals from religious orders (nuns, priests, brothers) from across the United States starting in 1994. MAP enrolled lay persons from across northeastern Illinois. Each study administers a battery of 21 cognitive performance tests annually of which 19 are in common. Alzheimer’s Disease status was determined by a computer algorithm based on cognitive test performance with a series of discrete clinical judgments made in series by a neuropsychologist and a clinician. Persons were categorized as no cognitive impairment (NCI) if diagnosed without dementia or mild cognitive impairment (MCI). Diagnoses of dementia and AD conform to standard definitions. A clinician reviewed all cases determined by this algorithm to render a diagnosis blinded to data collected in prior years. In addition to dementia, five other diagnoses were determined by this approach including stroke, cognitive impairment due to stroke, parkinsonism, Parkinson’s disease, and depression. Most other diagnoses are by self report. Upon death, a summary diagnosis is made by a neurologist blinded to the post-mortem assessment. A post-mortem neuropathologic evaluation is performed that includes a uniform structured assessment of AD pathology, cerebral infarcts, Lewy body disease, and other pathologies common in aging and dementia. The procedures follow those outlined by the pathologic dataset recommended by the National Alzheimer’s Disease Coordinating Center and pathologic diagnoses of AD use NIA-Reagan and modified CERAD criteria, and the staging of neurofibrillary pathology uses Braak Staging. Both studies are conducted by the same clinical and pathologic data collection teams with extensive item-level harmonization allowing the data to be efficiently merged.

### ROSMAP Genotype data Processing

Plink2 was used to perform operations on the genotype file (see link https://www.cog-genomics.org/plink/1.9/general_usage#cite), and positions were liftovered from hg18 to hg19 (http://genome.ucsc.edu/cgi-bin/hgLiftOver). Picard was used to sort the resulting genotype file, and variants with more than 2% missing values, minor allele frequency less than 1%, Hardy-Weinberg equilibrium less than 10E-6 as well as inbred samples (inbreeding coefficient >0.15) and samples with more than 2% missing values were removed using Plink2. Starting with 750,173 variants in 1,708 individuals for ROSMAP, 736,073 variants in 1,091 individuals for ROSMAP were left after quality control.

### ROSMAP RNAseq data processing

The RNA-seq BAM files were sorted using samtools[17] and converted to fastq files using the SamToFastq function (Picard 1.138, http://broadinstitute.github.io/picard/). RAPiD[18] was used to generate a count matrix for the gene expression data and generate a vcf file for each sample aligned to hg19 from the fastq files.

### Imputation

After quality control, we used 1000 Genomes data[19] and IMPUTEv2[20] to impute untyped variants. Imputed variants were removed if they failed any of the previously listed quality control criteria or had information scores < 0.6. After imputation we had 7,132,687 variants in Mayo and 9,333,139 variants in ROSMAP.

### De-convolute RNA-seq data into Microglia-specific Expression Residual

ROSMAP RNA-seq read count expression data was normalized using log2 counts per million (CPM) and the TMM method[21] implemented in edgeR[22]. Genes with over 1 CPM in at least 30% of the experiments were retained. We then used precision weights as implemented in the voom function from the limma[23] R package to further normalize the gene counts. MAYO expression data was normalized using the CQN R package (see above). For ROSMAP, expression residuals were obtained by correcting for the effects of technical (study, sequencing batch), sample-specific (post-mortem interval (PMI), RNA integrity number (RIN), exonic mapping rate) and patient-specific covariates (sex, educational attainment, age at death). For Mayo we adjusted for a slightly different set of covariates due to availability of recorded measurements (source of sample, sequencing batch, RIN, exonic mapping rate, sex, age at death). For both ROSMAP and MAYO data, we computed the exonic mapping rate using RNAseQC[24]. The exonic mapping rate was also included in the covariates. Adjustment for covariates was done using the limma R package.

Further, and performed together with the above listed covariate adjustments, for both ROSMAP and Mayo, we also adjusted for 5 cell type markers[25]: ENO2 [neuron], CD68 [microglial], CD34 [endothelial], OLIG2 [oligodendrocyte], GFAP [astrocyte]. To obtain expression residuals that mimic expression patterns seen in microglial cells, we added, for every gene, the CD68 effects estimated by the linear regression models back to the expression residuals.

The final microglia-specific expression residual data available for analysis included 20,276 genes for 612 individuals (ROSMAP) and 19,885 genes for 266 individuals (Mayo), with 18,408 genes in common to the two datasets.

### Rationalize and Validate Single-gene Biomarker for Bulk-tissue RNA-seq De-convolution

#### Compare sc-RNAseq derived Biomarker Genes per Cell Type

We assembled cell-type specific biomarkers derived from existing single-cell RNAseq data for neurons, microglial, astrocyte, endothelial and oligodendrocyte respectively. These biomarkers are included in the lists (Supplementary File S6). In each cell type, we compared the biomarker genes from every pair of studies and calculated the significance of the overlap by Exact Fisher’s test. The FDR is used to correct for multiple testing.

#### PCA analysis of sc-RNAseq derived Biomarker Expression in AMP-AD Data

We first merged biomarker list from different scRNA-seq studies for each cell type. Then, extracted the gene expression matrix of merged biomarkers from the ROSMAP and MAYO RNA-seq data. Next, we applied principal component analysis (PCA) on the extracted RNA-seq sub-matrix.

#### Compare sc-RNAseq derived Biomarker Genes with AMP-AD AGORA Targets

We merged all biomarkers for each cell type and calculated the percentage of overlapping with AGORA Targets. To evaluate the significance of this overlap, we simulated a background distribution of overlap by randomly selected the same number of genes from background genes (taking the non-duplicate union of genes in MAYO and ROSMAP RNA-seq data) per cell type, and compare the randomly generated “pseudo” biomarker list to AMP-AD AGORA Targets to generate a overlapping percentage. We repeated the random simulation 10,000 times to construct the background distribution. The p-value is then calculated by comparing true percentage to the background distribution per cell type.

#### Evaluate Robustness of Single-gene Biomarker derived Microglial-specific Residuals to scRNAseq Biomarker derived Microglial-specific Residuals

By using PSEA, we estimated the variance component of the bulk-tissue RNAseq data in ROSMAP and MAYO dataset explained by our single-gene microglial biomarker (CD68). Next, we randomly select a subset of biomarkers of each cell type from the scRNAseq-derived biomarkers (Supplementary File S6), then applied PSEA again to estimate the variance component of the bulk-tissue RNAseq data explained by the simulated subset of biomarkers. Then, we calculated the Pearson correlation of each gene between our single-gene microglial residuals with the simulated microglial residuals. We repeated this procedure 1,000 times to construct a distribution of the correlations.

Next, we intend to construct a background distribution of correlation. To this end, we again randomly select a subset of “pseudo” biomarkers of each cell type from the background genes (see above), then applied PSEA to estimate the variance component of the bulk-tissue RNAseq data explained by the simulated “pseudo” biomarkers. Then, we calculated the Pearson correlation of each gene between our single-gene microglial residuals with the simulated “pseudo” microglial residuals. We repeated this procedure 1,000 times to construct a distribution of the correlations. Lastly, we applied t-test to calculate the p-value based on the two distributions.

### Computational Analysis of Microglial-specific Gene Expression Data eQTL analysis

Expression quantitative trait loci (eQTL) analysis was performed using the R package MatrixEQTL v2.1.1[26] using QCed genotypes and normalized and covariate-adjusted celltype-specific expression residuals. cis-eQTL analysis considered markers within 1Mb of the transcription state site of each gene. False discovery rates were computed using the Benjamini–Hochberg procedure[27].

#### Differential Expression (DE) Analysis

We interrogated the celltype-specific residual expression data for genes differentially expressed between AD cases and healthy controls using linear models, as implemented in the R package limma[23]. Significance was assessed using Benjamini-Hochberg corrected p-values < 5%.

#### Co-expression networks Analysis

Co-expression networks were constructed using the coexpp R package[28] (Michael Linderman and Bin Zhang (2011). coexpp: Large-scale Co-expression network creation and manipulation using WGCNA. R package version 0.1.0. https://bitbucket.org/multiscale/coexpp). A soft thresholding parameter value of 6.5 is used to power the expression correlations. Seeding gene lists for the predictive networks were obtained by selecting genes in co-expression modules that were statistically enriched (FDR adjusted p-value < 0.05) for DE genes or astrocyte or microglial cell markers (lists of the latter two were obtained from [29]).

### Key Driver Analysis

To do Key Driver Analysis, we used the R package KDA[30] (KDA R package version 0.1, available at http://research.mssm.edu/multiscalenetwork/Resources.html). The package first defines a background sub-network by looking for a neighborhood K-step away from each node in the target gene list in the network. Then, stemming from each node in this sub-network, it assesses the enrichment in its k-step (k varies from 1 to K) downstream neighborhood for the target gene list. In this analysis, we used K = 6.

### Predictive networks Modeling and In-silico Prediction Validation

Though the co-expression network modules capture highly co-regulated genes operating in coherent biological pathways, these modules do not reflect the probabilistic causal information needed to identify key driver genes. Conventional Bayesian networks (BN) have been widely used to infer causal structures among genes given gene expression data, however, BN has significant limitations when it comes to infer opposite causality given the symmetry of joint probability. Recent work[31] has demonstrated that the bottom-up causality inference can accurately distinguish true causality from opposite causality in equivalent class. In this study, we developed a novel computational network model, called Predictive Network, by integrating conventional (top-down) Bayesian network with the bottom-up causality inference to address the problem of opposite causality inference in BN. Our causal predictive network pipeline incorporated multi-scale omics data, such as genotypes and transcriptomic profiles, in ROSMAP and MAYO dataset (de-convoluted microglial-specific residuals) to build causal predictive networks separately in ROSMAP and MAYO.

The predictive network model captures causal regulations among genes, which allows us to generate (in-silico) predictions upon perturbations, e.g. shRNA. Previously[32], we developed an integrative method, called Qualitative-constrained Maximal-a-Posterior (QMAP), to estimate the parameters of probabilistic graphical models. This method has been demonstrated to outperform traditional Maximal-a-Posterior (MAP) estimation without prior information. In this paper, we extended QMAP to integrate infinite number of resources of (priori) information on genetic regulations and big training data, to estimate parameters for constructed MAYO- and ROSMAP-microglial networks. Firstly, we collected multiple resources of prior information: i) we checked each edge in the MAYO- and ROSMAP-microglial network model against pathway knowledgebase, such as CPDB and String; ii) we applied linear regression to the (continuous) residual data to estimate the interaction type of each edge; Secondly, we integrated the two resources of prior with data to derive the parameters.

To predict gene expression fold-change upon shRNA against each key driver (HCK, FCER1G, LAPTM5), we developed three-step generalization procedure. First, we extracted the total sub-network of three key drivers from the constructed MAYO- and ROSMAP-microglial predictive networks (MAYO-sub, ROSMAP-sub) and included the Markov blanket of all top nodes in these sub-networks from the original MAYO-/ROSMAP-microglial networks. Second, we simulated the predictive network under wild-type (unperturbed) AD condition where probability of every top node is initialized according to the MAYO-/ROSMAP-microglial residual data under AD condition. Thirdly, we simulated the predictive network by perturbing key drivers under AD condition where probability of each key driver is initialized according to its knockdown level measured by Tagman (Figure S9) and other top nodes are initialized according to wild-type AD condition.

To calculate simulated gene expression fold-change, we used previously developed method [33, 34] to calculate the ratio of marginal probability of 18 measured target genes and compared to the experimental gene expression fold-change by Pearson correlation.

### Induction of Monocyte-Derived Microglia-like Cells (MDMi)

Peripheral blood mononuclear cells (PBMCs) were separated by Lymphoprep gradient centrifugation (StemCell Technologies). PBMCs were frozen at a concentration of 1–3 × 10^7^ cells ml^−1^ in 10% DMSO (Sigma-Aldrich)/90% fetal bovine serum (vol/vol, Corning). Prior to each study, aliquots of frozen PBMCs from the PhenoGenetic cohort were thawed and washed in 10 ml PBS. Monocytes were positively selected from whole PBMCs using anti-CD14+ microbeads (Miltenyi Biotech) and plated at the following densities per well: 1 x 10^5^ cells (96-well plate). To induce the differentiation of MDMi, monocytes were incubated in serum-free conditions using RPMI-1640 Glutamax (Life Technologies) with 1% penicillin/streptomycin (Lonza) and 2.5 μg/ml Fungizone (Life Technologies) and a mixture of the following human recombinant cytokines: M-CSF (10 ng/ml; Biolegend 574806), GM-CSF (10 ng/ml; R&D Systems 215-GM-010/CF), NGF-β (10 ng/ml; R&D Systems 256-GF-100), CCL2 (100 ng/ml; Biolegend 571404) at standard humidified culture conditions (37°C, 5% CO_2_) for up to 10 days[35].

### Lentivirus mediated shRNA triggered knockdown in primary monocyte derived microglia like cells (MDMi)

*shRNA lentiviral particle preparation:* Vpx viral particles were made using 293 T cells. On day 1, 293T cells were transfected using Lipofectamine 2000 (Thermo Fisher Scientific) along with envelope and packaging plasmids (Siv3+, pHEF VsVg a concentration of 1 μg/ml). On day 2, the culture medium was replaced by RPMI (Invitrogen) with 1% pennstrep and 1% fungizone (Amphotericin B) medium; the lentiviruses containing the Vpx particles were harvested 48 hours later and centrifuged at 400 *g* for 5 minutes at 4°C. The final product was filter sterilized using a 0.45-μm syringe filter (EMD Millipore). shRNA for target gene containing Lentiviral particles for each gene were obtained from the Broad Institute GPP (HCK, TRCN00000379914 and TRCN00000379408), FcER1G (TRCN0000057455 and TRCN0000057457) and LAPTM5 (TRCN0000428031 and TRCN0000429201).

*Lentivirus mediated knockdown of MDMi:* The monocytes were isolated from 9 healthy subjects and differentiated to MDMi using the above-mentioned protocol and plated on 96 well - temperature sensitive plate (Life Technologies #) as well as regular 384 well plates. On day 4, in the process of differentiation, Media was changed and lentivirus containing the Vpx particles and the lentivirus containing the shRNA for target gene from Broad were added to the cells, MDMi were maintained in the RPMI media with MDMi cocktail. On day 7, the transduced cells were selected using puromycin (Thermofisher scientific) at a concentration of (3ug/ml) conc. On day 10, the cells were lysed and gene expression assays (qPCR) was performed to validate expression of HCK, FcER1G and LAPTM5 as well as key downstream genes for each.

To assess statistical significance of differences in gene expression of knocked-down genes or genes downstream of these, we made use of linear mixed models, accounting for the multiple technical replicates for each biological replicate via random intercepts. These models also allowed us to deal with the fact that the experimental design was unbalanced with only two technical replicates for the empty control and three technical replicates for each biological replicates using shRNA triggered constructs.

#### Quantitative Real Time-Polymerase Chain Reaction (qRT-PCR)

RNA was extracted from each sample using RNeasy micro kit (Qiagen, USA). Genomic DNA contamination was minimized by spinning samples using a genomic DNA column (gDNA) according to the manufacturer’s instructions. RNA was reverse transcribed into cDNA using a Taqman Reverse Transcription kit (Invitrogen). qPCR was performed using TaqMan® Fast Advanced Master Mix (Applied Biosystems) and run on a Light cycler 480 System (Roche, USA). The cycling conditions consisted of 90 °C for 10 min and 40 cycles of 95 °C for 20 s followed by 60 °C for 30 sec. Samples were assayed with 2 technical replicates. mRNA levels were normalized relative to B2M by the formula 2^−ΔCt^, where ΔCt = CtmRNA-X – CtB2M.

### A**β**1-42 Uptake Assay

We tested the uptake ability of lentivirus mediated shRNA triggered downregulated HCK and FcER1G MDMi using HiLyte™ Fluor 488-labeled beta amyloid 1-42(Anaspec AS-60479-01) for a period of 10 days in RPMI media with MDMi cocktail (Katie’s paper). On day 10, the media was replaced with media containing 1.5ug/ml of HiLyte™ Fluor 488-labeled beta amyloid 1-42 for 2 h at 37 °C. After 2 hours, cells were washed three times with PBS and fixed in 4% PFA for 15 min. The cells were then imaged using confocal image express C (Harvard, Longwood ICCB). The mean fluorescence intensity was measured using the Multi-wavelength scoring program. Data shown is the mean fluorescence intensity for each subject. Two-tailed, paired t-tests were used to determine statistical significance.

---

**Figure S1.**
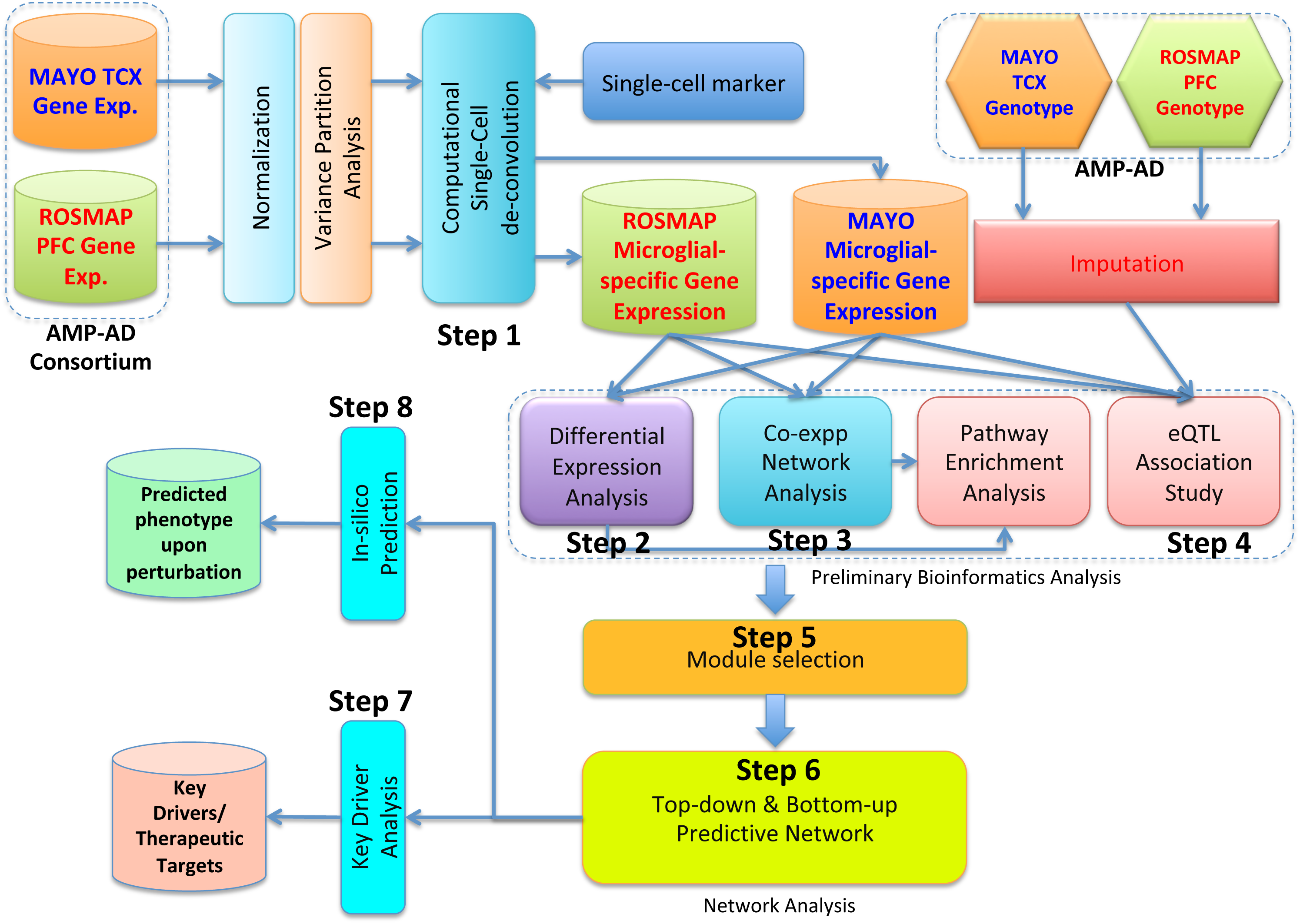

**Figure S2.**
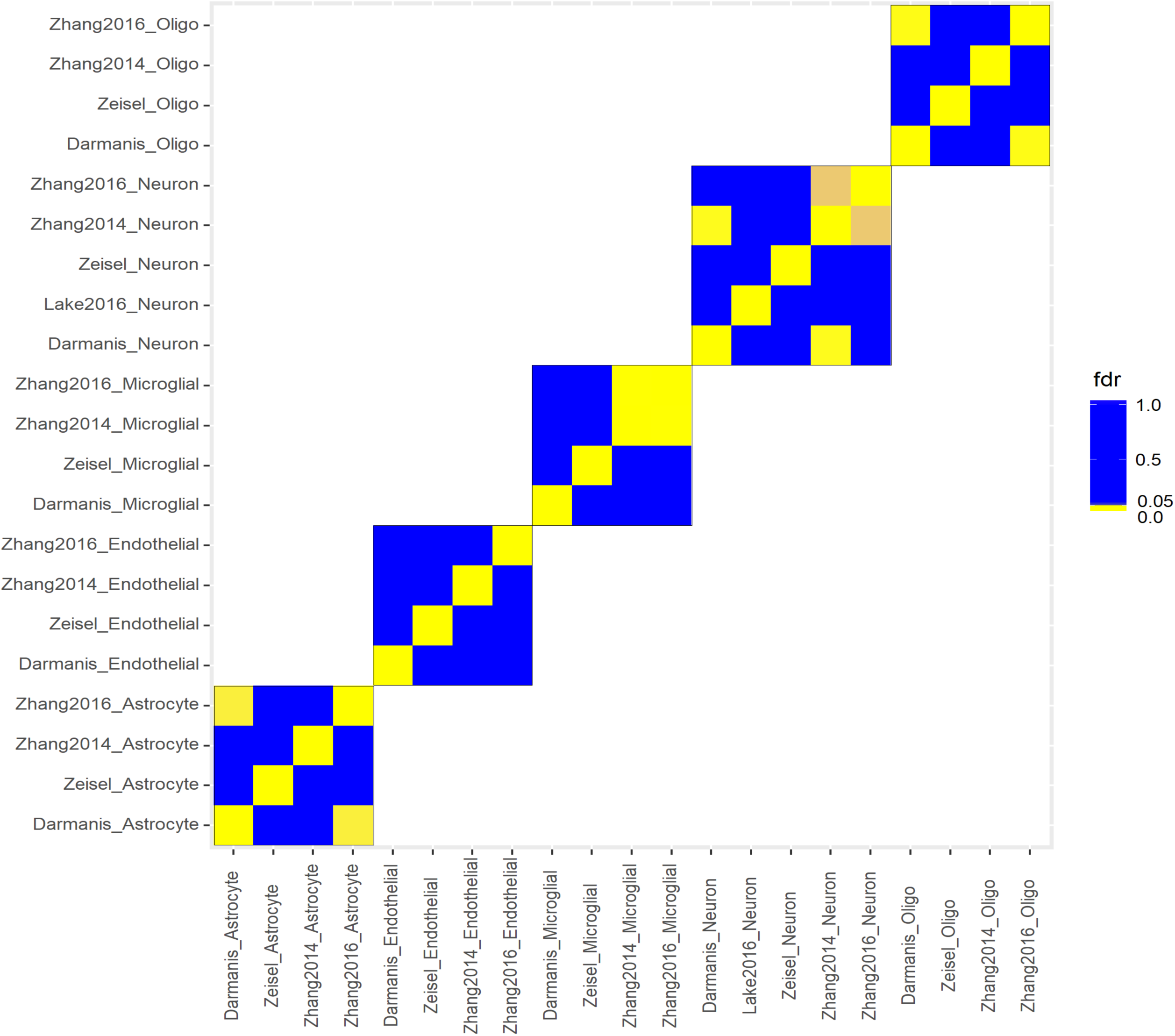

**Figure S3.**
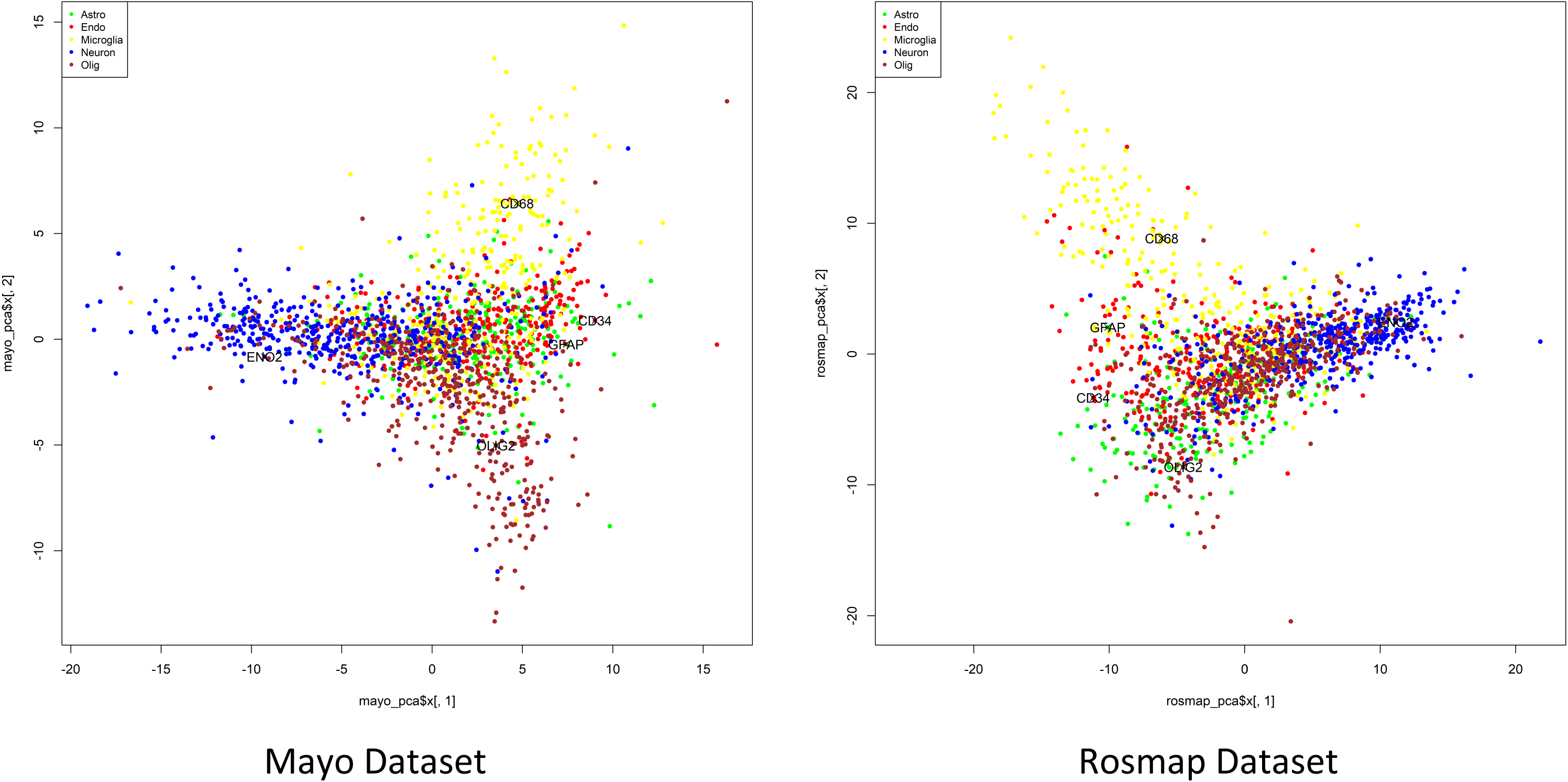

**Figure S4.**
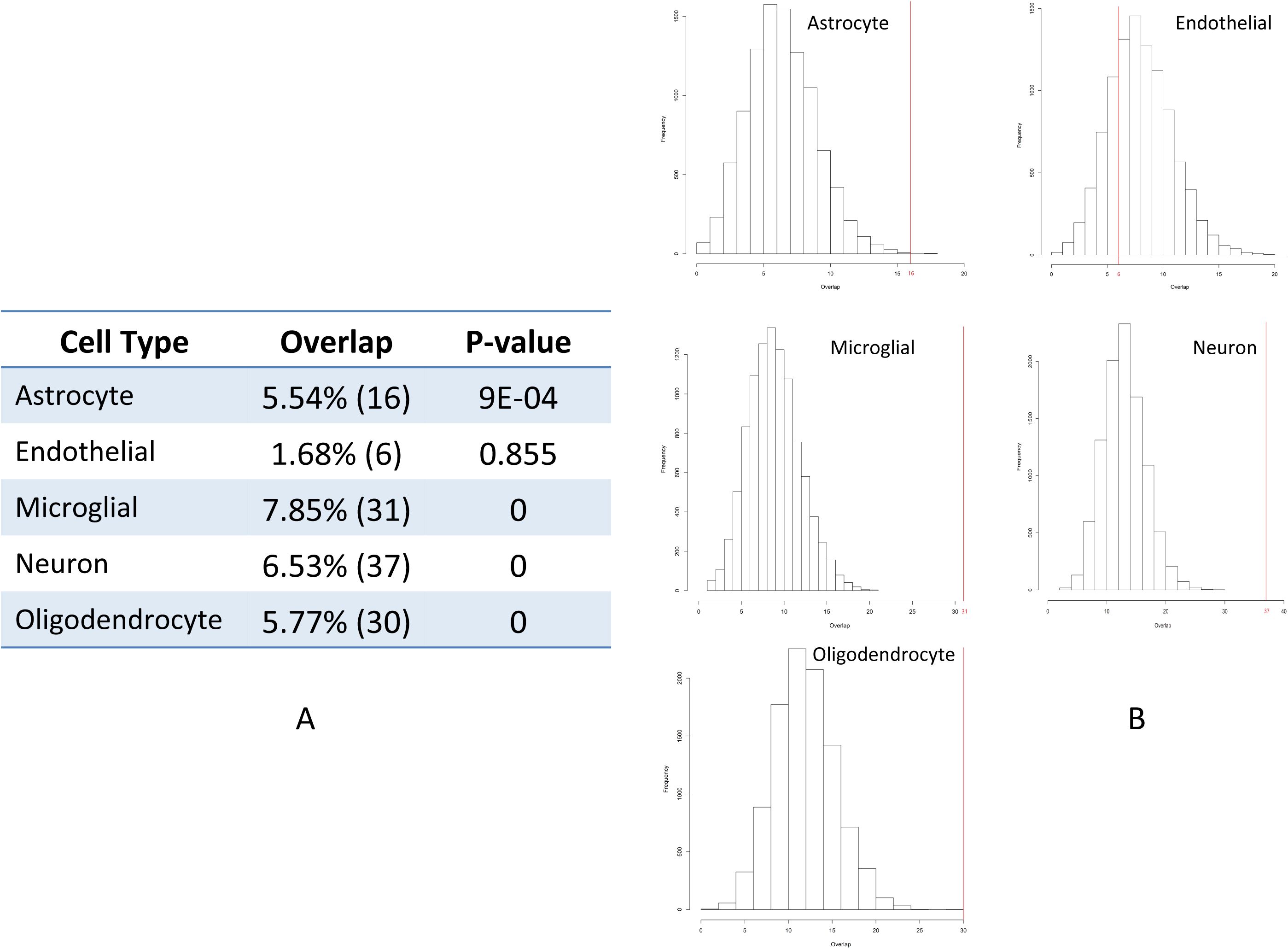

**Figure S5.**
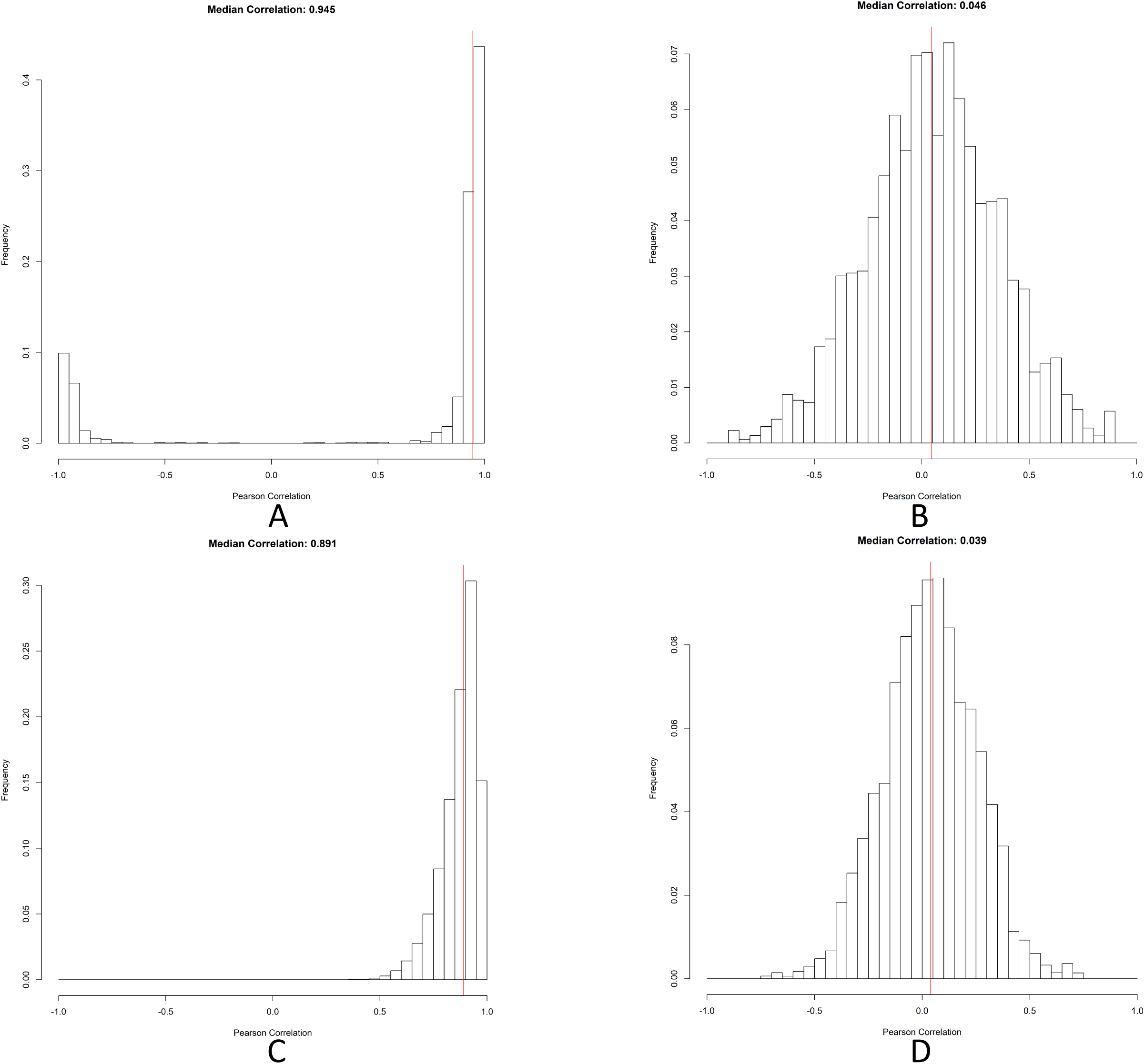

**Figure S6.**
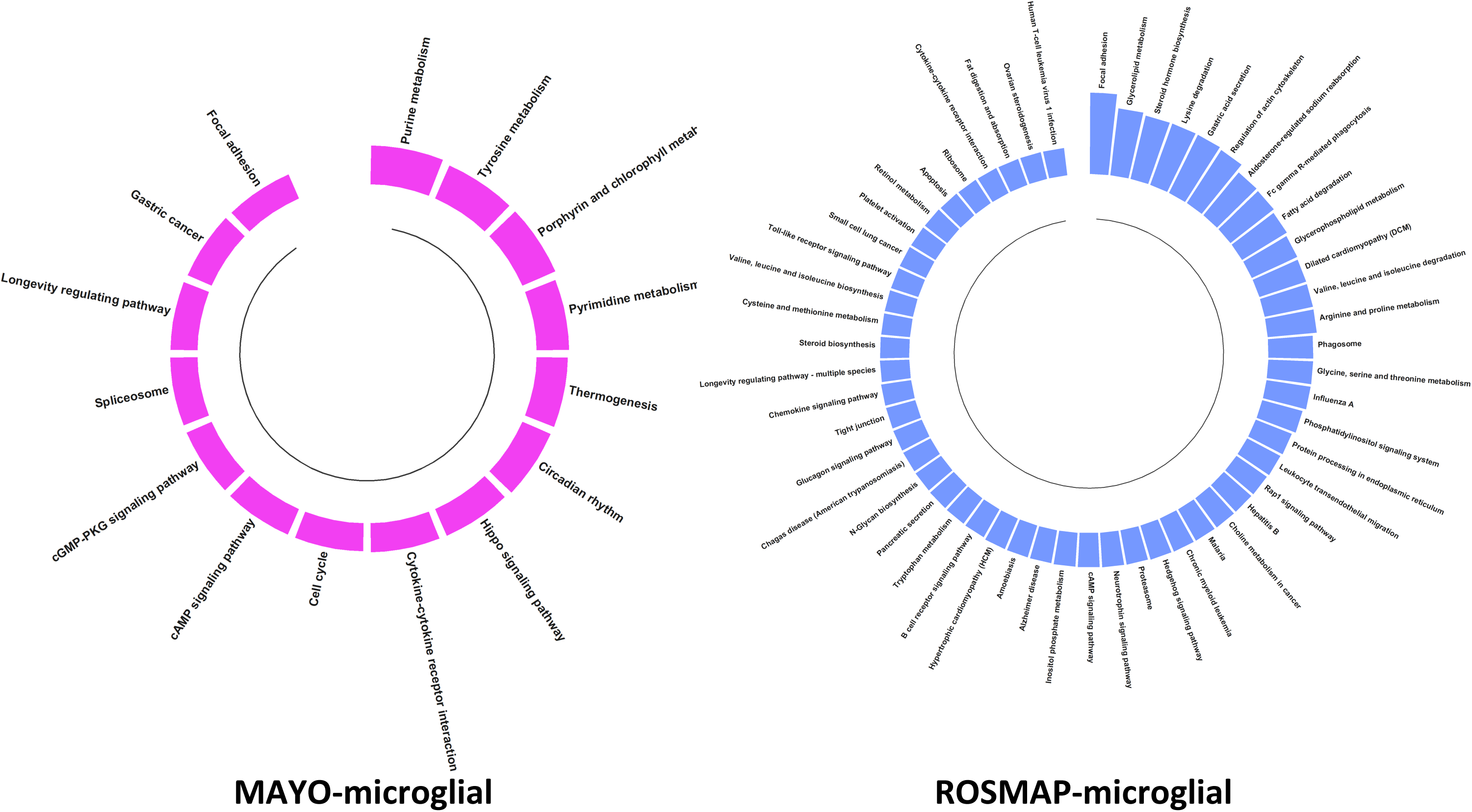

**Figure S7.**
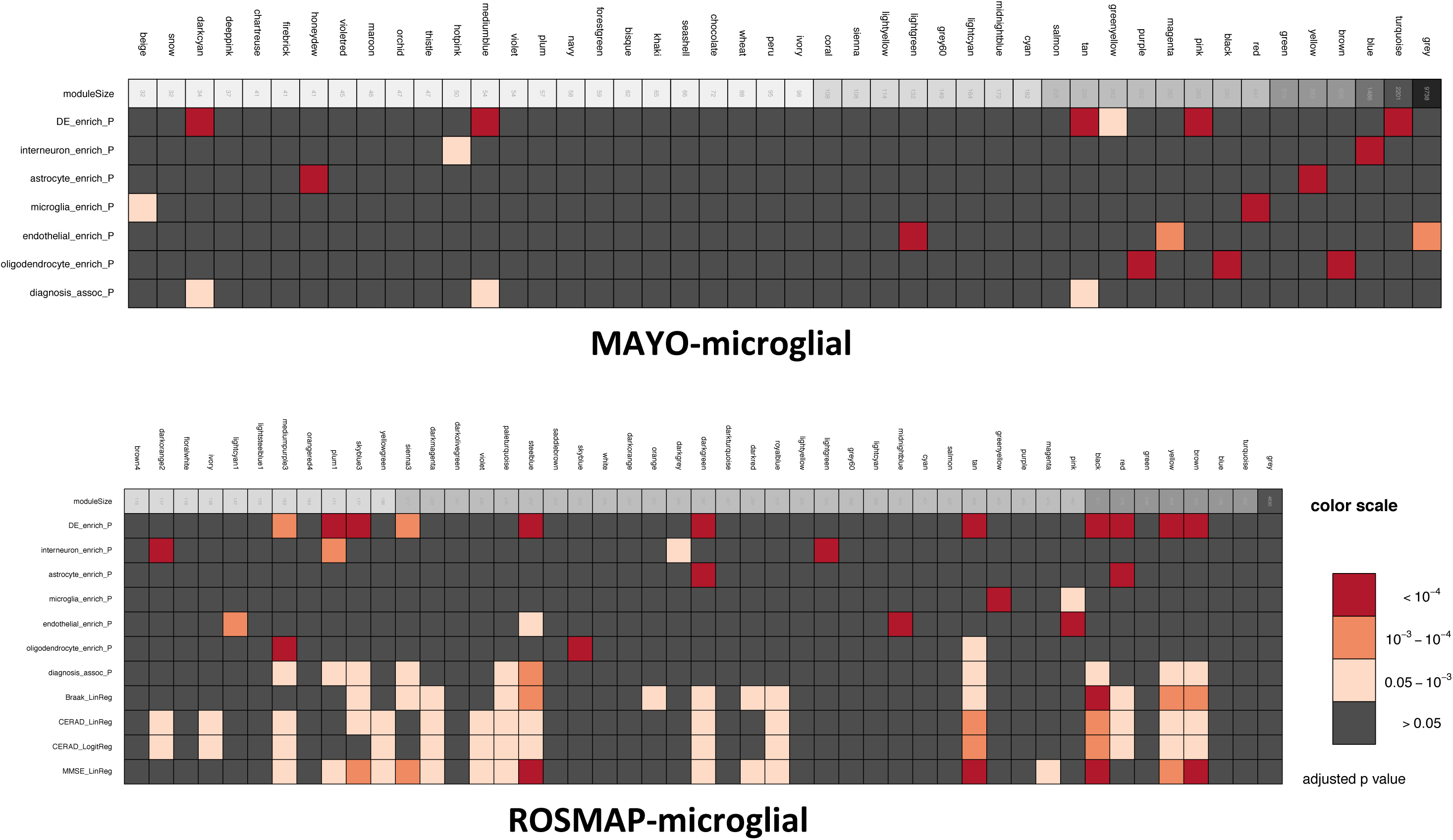

**Figure S8.**
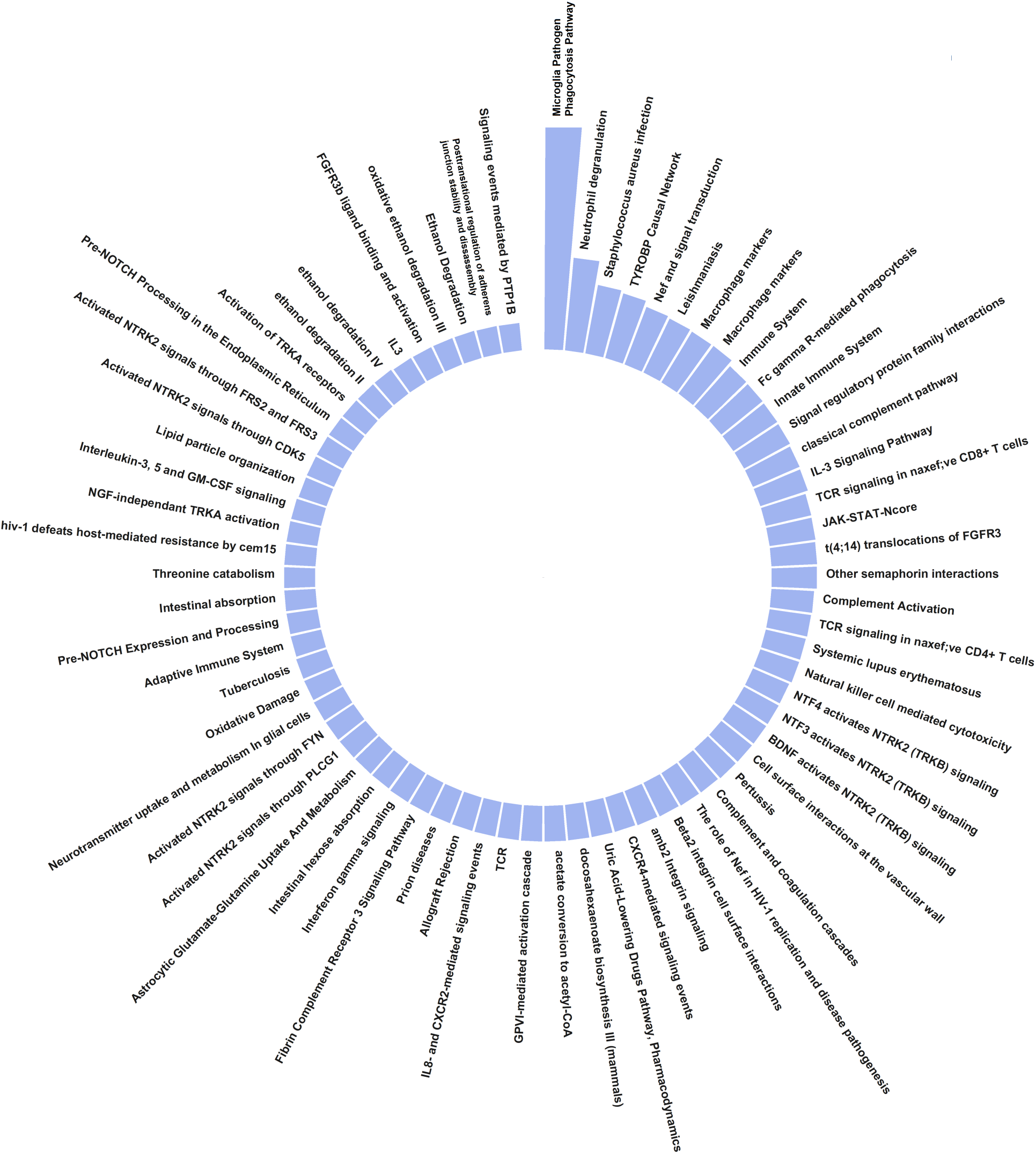

**Figure S9.**
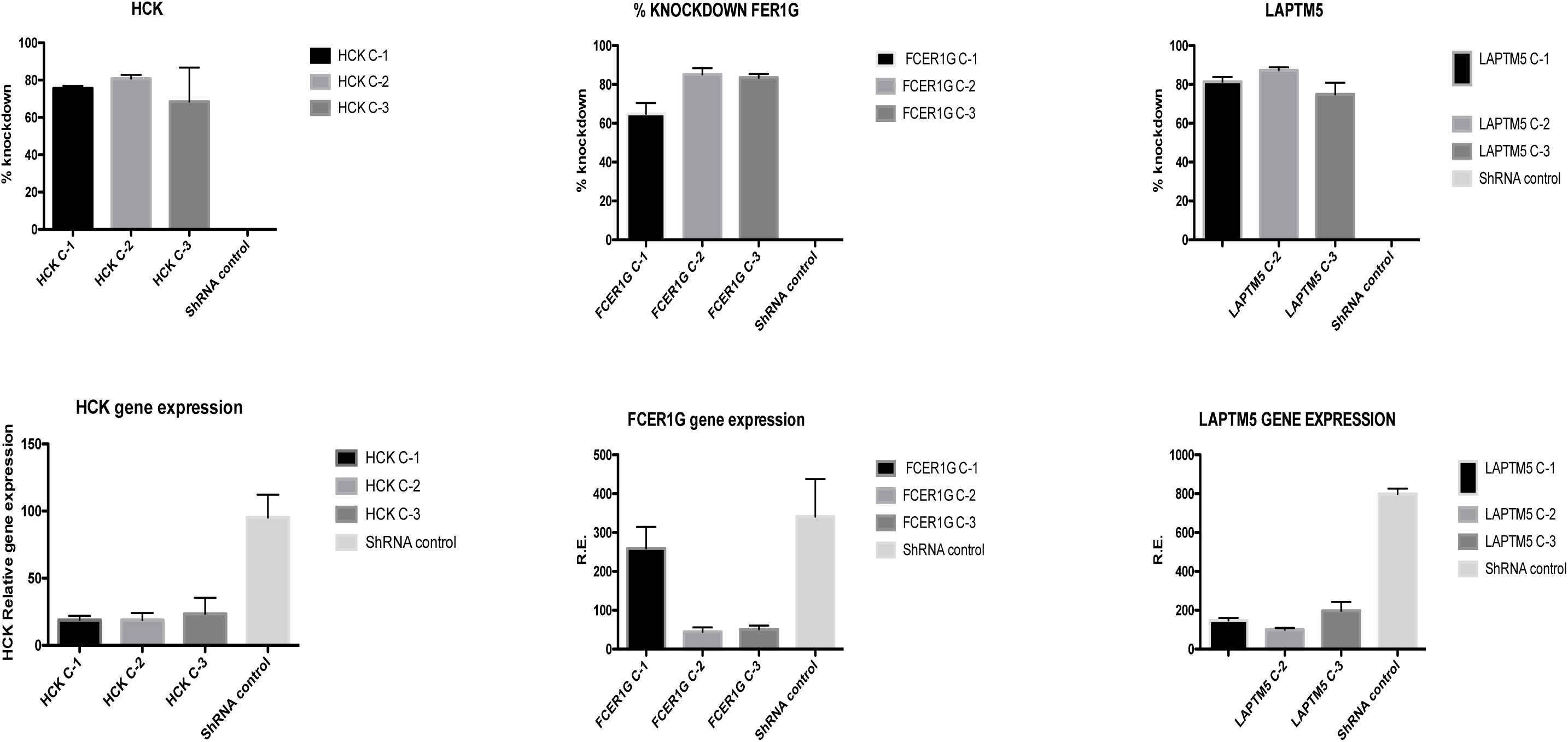

